# Structural and biochemical characterization of the ALX4 dimer reveals novel insights into how disease alleles impact ALX4 function

**DOI:** 10.1101/2024.09.10.612331

**Authors:** Brittany Cain, Zhenyu Yuan, Evelyn Thoman, Rhett A. Kovall, Brian Gebelein

## Abstract

How homeodomain proteins gain sufficient DNA binding specificity to regulate diverse processes has been a long-standing question. Here, we determine how the ALX4 Paired-like protein achieves DNA binding specificity for a TAAT – NNN – ATTA dimer site. We first show that ALX4 binds this motif independently of its co-factor, TWIST1, in cranial neural crest cells. Structural analysis identified seven ALX4 residues that participate in dimer binding, many of which are conserved across the Paired-like family, but not other homeodomain proteins. Unexpectedly, the two ALX4 proteins within the dimer use distinct residues to form asymmetric protein-protein and protein-DNA interactions to mediate cooperativity. Moreover, we found that ALX4 cooperativity is required for transcriptional activation and that ALX4 disease variants cause distinct molecular defects that include loss of cooperativity. These findings provide new insights into how Paired-like factors gain DNA specificity and show how disease variants can be stratified based on their molecular defects.

## 2. Introduction

The homeodomain (HD) family is one of the largest families of transcription factors (TFs), consisting of ∼200 proteins in humans that are classified by their highly conserved helix-turn-helix DNA binding domain ^1^. *In vitro* DNA binding assays have shown that HDs largely bind highly similar AT-rich DNA sequences ^2–4^, primarily through contacts with the N-terminal Arginine Rich Motif (ARM) and the third helix, which is also known as the recognition helix ^5^. However, the mechanisms by which these HDs achieve sufficient DNA binding specificity to regulate their distinct targets *in vivo* are not well understood.

Several mechanisms have been described to address this paradox. First, some HD subfamilies such as the Paired, Prospero, and CUT classes encode additional DNA binding domains that bind distinct sequences. Second, studies have shown that while consensus high affinity sites are often bound by many HD factors in a relatively indiscriminate manner, low affinity sites are typically bound by fewer HD factors and thereby result in enhanced target specificity ^6–8^. Third, some HD proteins form complexes that facilitate binding to distinct and/or longer recognition sequences. For example, the HOX factors form complexes with the PBX and MEIS HD proteins to increase DNA binding specificity ^9^. Additionally, members of the Paired-like family, which share sequence conservation with the HD encoded by Paired (PAX) family members but lack the accompanying Paired DNA binding domain (Bürglin & Affolter, 2016), have been shown to form dimers that both increase DNA binding specificity ^11,12^ and alter transcriptional output ^11,13^. Moreover, we previously used a computational pipeline to predict cooperative homodimer binding of HD TFs from high throughput systematic evolution of ligands by exponential enrichment (HT-SELEX) data and found that several, but not all, Paired-like members cooperatively bind a TAAT – NNN – ATTA (P3) dimer site ^14^. Currently, however, it is not well understood which HD residues within the Paired-like factors confer cooperative DNA binding to these P3 sites.

The mechanisms by which Paired-like factors bind cooperatively have been previously extrapolated from a crystal structure of a modified Paired (Prd) *Drosophila* HD bound to DNA ^15^. However, there are several caveats to using this structure as a model. First, Prd is a PAX TF, containing a Paired domain and a HD that both possess the capability to bind DNA, and Prd preferentially uses these domains to bind a distinct hybrid site that does not resemble a HD site ^16,17^. Second, the HD of the wild type Prd protein, which contains a serine at position 50 (S50) within the third helix, binds cooperatively with equal affinity to a TAAT – NN – ATTA sequence (P2 site) and the P3 site, whereas the Paired-like factors encode a Q50 HD and strongly prefer binding the P3 site ^12^. Hence, the Prd HD only favors the P3 site when a S50Q mutation is created ^12^, making it unclear whether the Paired-like factors use similar interactions and mechanisms to bind cooperatively.

Here, we focus on defining the mechanisms used by the Paired-like factor, ALX4, to cooperatively bind DNA. Genetic studies in mice and humans have shown that ALX4 is critical for proper craniofacial development. Alx4 null mutations in mice cause enlarged parietal foramina in which the parietal bones fail to fuse resulting in a gap at the sagittal sinus of the skull as well as polydactyly and loss of cartilage in the limbs ^18,19^. Consistent with these phenotypes, autosomal dominant and recessive ALX4 variants in humans have been associated with a variety of craniofacial defects. Patients with a homozygous nonsense variant display frontonasal abnormalities, alopecia, and sagittal suture defects ^20^, and missense variants in ALX4 are implicated in three distinct developmental disorders: Enlarged parietal foramina ^21–25^; sagittal suture craniosynostosis in which the parietal bones fuse too early in development leading to stunted brain growth, increased intracranial pressure, and scaphocephaly ^26^; and frontonasal dysplasia in which patients exhibit abnormal development of craniofacial features ^21,25,27^. The molecular mechanisms behind how many of these ALX4 missense variants cause these distinct disease phenotypes are not well understood.

A recent study found that ALX4 can also cooperatively bind DNA with the basic helix loop helix (bHLH) factor, TWIST1, to a composite site called the coordinator that contains a bHLH E-box site and a HD site spaced 6bp apart ^28^. This motif was originally identified due to it being highly enriched in human specific craniofacial enhancers compared to chimp, suggesting that the coordinator motif and the TFs that bind to it drive human craniofacial development ^29^. To identify the TFs that bind the coordinator site, Kim et al generated a large amount of genomic binding data in human cranial neural crest cells (hCNCCs) to show that TWIST1 binds to the E-box sequence, drives chromatin opening, promotes HD recruitment of either ALX1, ALX4, MSX1, or PRRX1, and promotes enhancer acetylation at the coordinator motif ^28^. Taken together, these studies suggest that ALX4 uses multiple modes of DNA binding to regulate the expression of target genes required for craniofacial development.

In this study, we use a combination of structural, biochemical, and bioinformatic approaches to both determine how ALX4 homodimers cooperatively bind DNA and determine how disease variants within the HD impact ALX4’s molecular functions. First, we took advantage of published ALX4 genomic binding data from differentiated hCNCCs to identify diverse ALX4 bound site types ^28^. This analysis revealed ALX4 binding to the TAAT – NNN – ATTA (P3) site was largely independent of TWIST1, whereas ALX4 binding to the monomer and coordinator sites was largely TWIST1-dependent. Next, we used crystallography to solve the ALX4 HD dimer structure bound to a P3 site and found that while it resembles the Prd S50Q HD dimer ^15^, ALX4 forms unique asymmetric interactions between the two bound ALX4 proteins. Based on this structure, we used mutagenesis studies to define residues critical for cooperative DNA binding and assessed how five ALX4 missense variants associated with human birth defects impact DNA binding, cooperativity, and transcriptional output. Importantly, disease variants could be stratified based on their molecular impact which includes complete loss of DNA binding, loss of cooperative DNA binding, loss of protein stability, and subcellular mis-localization, each of which leads to decreased transcriptional output on the P3 site. Overall, these studies reveal novel mechanisms by which ALX4 binds cooperatively to its P3 site and begin to define the gene regulatory impact of ALX4 cooperativity and its role in ALX4-associated disease.

## 3. Results

### Genomic binding of ALX4 to the P3 dimer site is independent of co-factor TWIST1

To identify the genomic DNA binding sites of ALX4, we utilized available CUT&RUN (C&R) data from human embryonic stem cells lines differentiated into hCNCCs ^28^. These cells contain an engineered TWIST1 locus that expresses a FKBP12^F36V^-tagged TWIST1 protein that undergoes rapid degradation following treatment with a dTAG-1 ligand, providing temporally controlled expression of TWIST1 (Figure 1A). With this system, Kim et al assessed ALX4 genomic binding in the presence and absence of TWIST1 ^28^. Since Kim et al primarily analyzed this data from the perspective of TWIST1 binding (i.e. focused on co-binding to TWIST1 bound regions), we reanalyzed the ALX4 C&R data to determine if ALX4 bound to TWIST1-independent loci as well as TWIST1-dependent loci. From this analysis, we identified 3,250 ALX4 bound regions in the genome (Figure 1B). Consistent with these regions containing active enhancers, ALX4 bound regions were highly accessible and had high signal for H3K27 acetylation (Supplementary Figure 1). After a 1 hour treatment with the dTAG-1 ligand that results in rapid degradation of the FKBP12^F36V^-tagged TWIST1 protein (Figure 1A), ALX4 no longer bound to 2867 of these regions, indicating that many ALX4 bound regions were dependent on TWIST1 (Figure 1B). However, 383 ALX4 genomic regions were bound similarly in the presence and absence of TWIST1.

**Figure 1.**
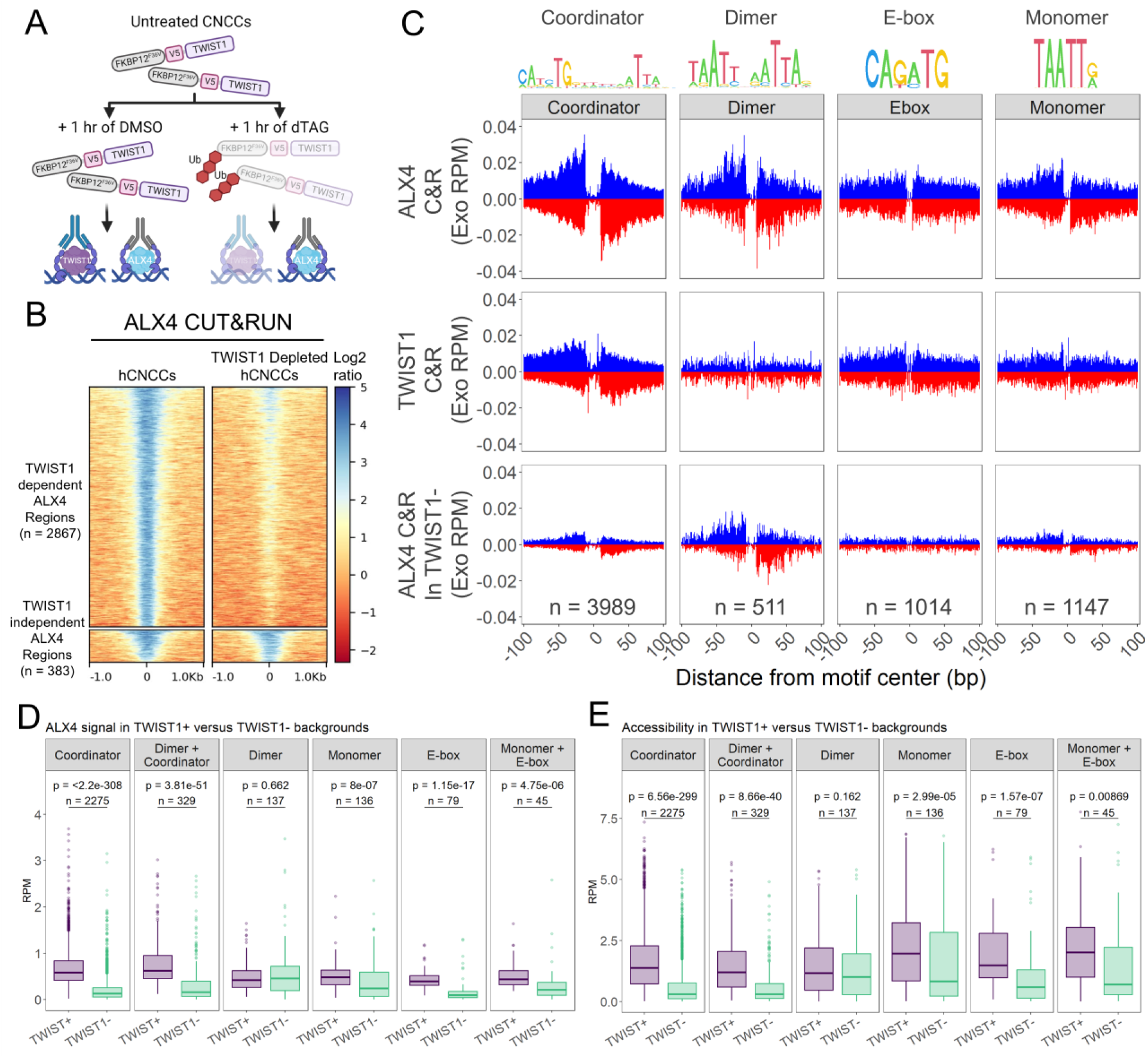
ALX4 genomic binding to the P3 dimer site is TWIST1 independent. **(A)** Schematics of experiments performed in Kim *et al* ^28^. **(B)** ALX4 bound to 3250 regions in wildtype conditions. 2867 (88%) regions were no longer bound in the dTAG treated samples in which TWIST1 was depleted (TWIST1 dependent ALX4 regions). 383 (12%) regions were maintained after TWIST1 depletion (TWIST1 independent ALX4 regions). The color bar denotes the log2 ratio of the immunoprecipitated sample versus the IgG control. **(C)** A Homer *de novo* motif analysis revealed four distinct site types that were enriched in the ALX4 C&R: the Coordinator ^28^, the P3 dimer site, an independent E-box, and an independent monomeric HD site. MNAse digestion patterns were used to discriminate between bound and unbound sites. TWIST1 dependence was assessed by footprinting the ALX4 bound sites with TWIST1 C&R and ALX4 C&R in a TWIST1 depleted background. **(D)** ALX4 binding signal decreases significantly after TWIST1 depletion for all peak sets except for peaks containing only the P3 dimer site. The 3250 ALX4 bound regions were categorized based on the presence of a coordinator site, a coordinator and dimer site, a dimer site, or only a monomer site or E-box or both independent sites. The average reads per million (RPM) after E. coli spike in normalization of the 200 bp window at the peak center were compared with a Wilcoxon rank sum test. **(E)** Chromatin accessibility decreases significantly after TWIST1 depletion for all peak sets except for peaks containing only the P3 dimer site. The average RPM of a 200 bp window at the peak centers were compared with a Wilcoxon rank sum test.

We next performed a *de novo* motif search ^30^ on the ALX4 bound regions and focused on four distinct binding motifs: the Coordinator, the P3 dimer site, an independent E-box site, and an independent HD monomer site (Figure 1C; Supplementary Figure 2). However, the presence of a motif does not guarantee TF binding, and the presence of a composite site does not guarantee that both TFs are bound at the same time. To stringently assess for TF binding, we first scanned ALX4 peaks for potential binding sites using Homer and then filtered potential binding sites by comparing MNase digestion patterns of the motif and nearby flanking windows. If a given site is bound, the motif should be protected from MNase cleavage, and the neighboring accessible sequences will be preferentially cleaved. To ensure sites were not scored in more than one category, we first identified all digest confirmed coordinator and P3 dimer sites and removed these prior to analyzing for independent E-box and HD monomer binding (Supplementary Figure 3A). Using this approach, we identified >500 bound sites for each site type within the 3,250 bound regions. Figure 1C displays the average 5’ MNase signal for the filtered site types, and the digestion signal of each site is shown in Supplementary Figure 3B. Note, the length of the protected region correlates with the length of the motif as expected.

The above data provides evidence that ALX4 binds four types of sites *in vivo*: Coordinator sites in complex with TWIST1, **<u>TAAT-</u>**NNN-**<u>ATTA</u>** (P3) dimer sites, independent E-box sites, and independent HD monomer sites. To assess which sites were also bound by TWIST1, we similarly analyzed the TWIST1 C&R for protected binding motifs. We found that TWIST1 was largely bound at the coordinator site, but not the three other site types (Figure 1C). Conversely, we wanted to identify ALX4 site types that are dependent versus independent of TWIST1. To do so, we performed footprint analysis for ALX4 bound sites in the TWIST1 depleted cells. Here, we found that the overall signal of ALX4 binding to the coordinator site was largely lost (Figure 1C). Note, the remaining weakened ALX4 footprint signal is consistent with the short-term treatment of dTAG being insufficient to deplete all TWIST1 at these binding sites. Interestingly, ALX4 P3 dimer site protection and signal were retained more so than other site types, suggesting that ALX4 P3 site binding is largely independent of TWIST1. In contrast, the footprint signal at monomer ALX4 binding was dramatically decreased. Taken together, these data suggest that ALX4 binding to the P3 dimer sites are largely independent of TWIST1, whereas ALX4 binding at monomer sites and coordinator sites are largely dependent on TWIST1.

To statistically analyze the biased retention of ALX4 P3 dimer site binding in the TWIST1 depleted background, we looked at the total binding signal and chromatin accessibility across peak centers. We first separated ALX4 peaks into six categories: (1) peaks that contained a coordinator site but not a P3 dimer site; (2) peaks that contained a coordinator and a P3 dimer site; (3) peaks that contained a P3 dimer site without a coordinator site; and (4 through 6) peaks that did not contain a coordinator site or a P3 dimer site but contained either a (4) ALX4 monomer site, (5) a TWIST1 E-box site, or (6) both sites. The majority of peaks contained coordinator sites, which emphasizes TWIST1’s strong influence on ALX4 binding ^28^. We then compared the ALX4 C&R binding signal as well as the ATAC-seq reads between wildtype cells and TWIST1 depleted cells. All peak categories significantly decreased in ALX4 binding signal (Figure 1D) and chromatin accessibility (Figure 1E) after TWIST1 depletion with the exception of peaks containing only P3 dimer sites, further demonstrating that the majority of ALX4 P3 dimer sites are bound independently of TWIST1. In contrast, the loss in accessibility in peaks containing only ALX4 monomer sites provides a potential explanation as to why these sites depend on TWIST1 regulation. Taken together, these data reveal that ALX4 genomic binding to the P3 dimer site, and not the other observed ALX4 binding sites, is TWIST1-independent.

### The ALX4 HD is sufficient to mediate cooperative DNA binding to the P3 site

We previously predicted and validated ALX4, as well as several other Paired-like HDs, form cooperative homodimers on a P3 site identified from HT-SELEX data that is identical to the TWIST1-independent ALX4 site found in the genomic analysis of hCNCCs ^14^. However, the mechanisms used by these HDs to bind DNA cooperatively as well as the regions of the HD required to facilitate cooperativity are largely unknown. Our prior studies revealed that an ALX4 protein containing the HD as well as relatively short peptide chains N- and C-terminal to the HD (aa 169-303) was highly cooperative on a probe containing a **<u>TAAT-</u>**TAG-**<u>ATTA</u>** P3 site ^14^. To assess if the ALX4 HD alone (aa 209-274) (Figure 2A) is sufficient to mediate cooperative DNA binding, we performed quantitative electrophoretic mobility shift assays (EMSAs) using the P3 site and a **<u>TAAT</u>** – TAGG – **<u>ATTA</u>** site (P4 site). Comparative analysis of ALX4 binding to the P3 and P4 sites revealed that the ALX4 HD is sufficient to cooperatively bind the palindromic sequence in a spacer-dependent manner (Figure 2B; Supplementary Figure 5A). We used Tau, which measures the multiplier by which the binding of the first ALX4 protein facilitates the binding of a second ALX4 protein, to quantify the cooperativity of ALX4 bound to a P3 site and P4 site ^12^; (See Methods for more information). A Tau greater than one demonstrates that binding of the first protein facilitates the binding of the second protein, a Tau equal to one demonstrates independent binding, and a Tau less than one demonstrates that the binding of the first protein hinders the binding of the second protein. On the P3 site, the Tau value (110±14) for the ALX4 HD revealed highly cooperative binding. In sharp contrast, when a single nucleotide is added between these two palindromic sites, cooperativity is lost as ALX4 fails to preferentially bind as a dimer complex (Tau = 1.5±1.4, Figure 2C). These findings demonstrate that the ALX4 HD is sufficient for cooperative DNA binding.

**Figure 2.**
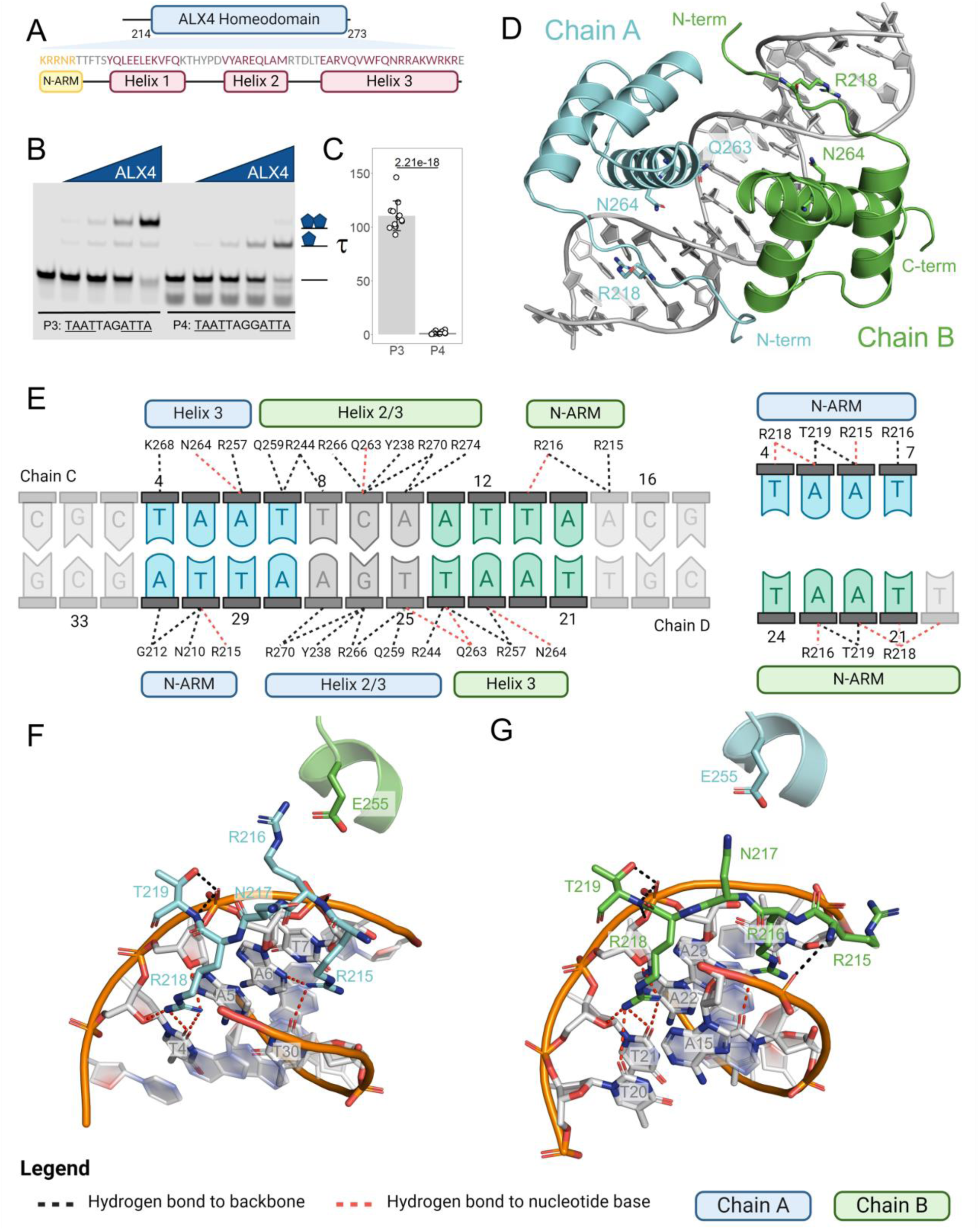
ALX4 cooperativity is DNA mediated. **(A)** A map of the ALX4 HD. **(B)** The ALX4 HD is sufficient to mediate cooperativity on the P3 site but unable to bind cooperatively to the P4 site. ALX4 protein (0, 37.5, 75, 150, and 300 nM) was combined with each respective fluorescent probe. A single replicate is shown here, and all three replicates are shown in Supplementary Figure 5A. Schematics of the protein-DNA complexes are shown to the right of the gel. **(C)** Tau cooperativity factors were calculated for every lane in which protein was added. Bars depict the average Tau for each probe and each dot represents a Tau from an independent binding reaction (n = 12). Error bars denote standard deviation. Tau factors were compared with an unpaired, two-sided student t-test. **(D)** X-ray structure of ALX4 reveals that ALX4 binds as a dimer to the P3 site in a head-to-head orientation. ALX4 Chains A and B are colored cyan and green, respectively; DNA is colored gray. ALX4 residues R218 (canonical HD numbering: R5), Q263 (Q50), and N264 (N51) are shown in a stick representation. **(E)** Schematic of ALX4-DNA interactions. Hydrogen bond interactions were determined by PDBePISA and required atoms to be within 4 Å. Chain A and B proteins are labeled in blue and green respectively. Hydrogen bond interactions to the DNA backbone are shown via black dotted lines whereas hydrogen bonds to nucleotide bases are shown in red. **(F-G)** Structural comparisons of protein-DNA interactions made by the ALX4 Chains A and B. **(F)** On Chain A, R215 (R2) inserts into the minor groove and forms key DNA contacts while R216 (R3) creates minimal contacts with DNA but instead forms a salt bridge with E255 (E42) of Chain B. **(G)** On Chain B, R216 (R3) inserts into the minor groove and forms key DNA contacts whereas R215 (R2) faces the loop between helix 1 and helix 2 of Chain A.

### ALX4 dimeric crystal structure reveals asymmetrical DNA contacts via the N-terminal ARM

To determine the structural basis for the mechanisms that underlie ALX4 cooperativity, we solved the crystal structure of the ALX4 HD (aa 209-274) bound to a P3 site (-CGC**<u>TAAT</u>**TCA**<u>ATTA</u>**ACG-) at 2.39 Å resolution (Figure 2D) as well as the ALX4 HD not bound to DNA at 2.39 Å resolution (Supplementary Figure 5B; Supplementary table 1). Both proteins in the dimer complex as well as the unbound protein share the same overall structure with the exception of the N-terminal ARM, which is largely unstructured off DNA and thereby differs in both conformation and electron density resolution (Supplementary Figure 5B). Here, we focus on the ALX4 dimer structure on DNA, where we label the two ALX4 proteins as Chain A (cyan) and Chain B (green), and separate our analysis of the ALX4 protein-to-DNA interactions (Figure 2E-G) and ALX4 protein-to-protein interactions (Figure 3).

**Figure 3.**
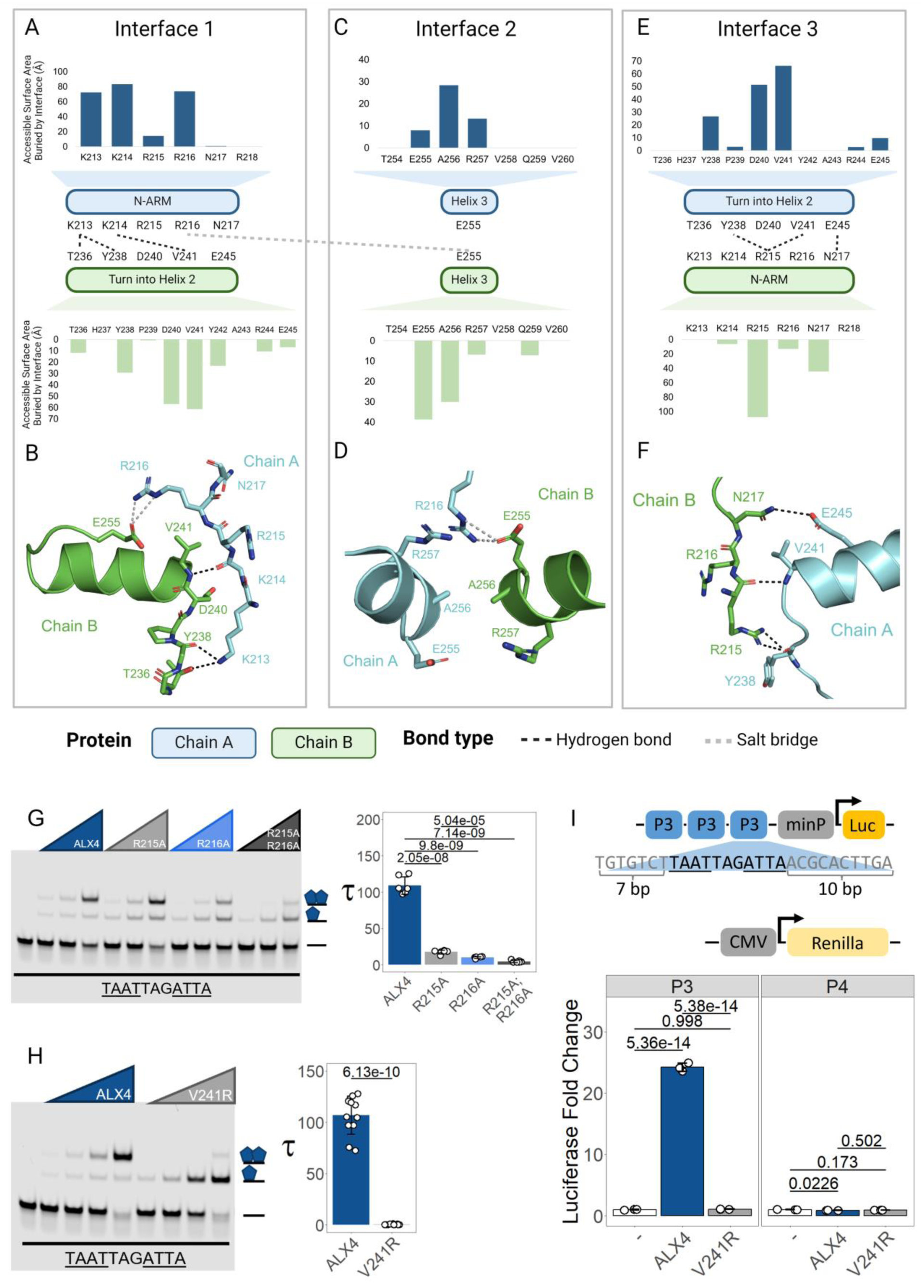
ALX4 cooperativity is facilitated by three protein-protein interfaces. **(A, C, E)** Van der waals interactions were calculated via the total accessible surface area of the residue that is buried by the other protein chain and are shown in the bar graphs. Polar contacts are shown via dotted lines between interacting residues: hydrogen bond contacts are shown with black dotted lines, whereas salt bridge contacts are shown via gray dotted lines. **(B, D, F)** Protein structure views of the protein-protein interfaces. **(A-B)** The N-terminal ARM (N-ARM) of Chain A contacts the turn between helix 1 and 2 as well as helix 3 of Chain B. **(B-C)** The start of both recognition helices forms a symmetrical interface. **(D-E)** The N-ARM of Chain B forms contacts with the turn between helix 1 and 2 of Chain A. **(G)** R215A (R2A), R216A (R3A), and R215A; R216A (R2A; R3A) mutations negatively affect ALX4 cooperativity in a compounding fashion. ALX4 protein (0, 50, 100, and 200 nM) were tested on the P3 fluorescent probe. Schematics of the protein-DNA complexes are shown to the right of the gel. Tau cooperativity factors were calculated for every lane in which protein was added. Bars depict the average Tau for each protein and each dot represents a Tau from an independent binding reaction. Error bars denote standard deviation. Tau factors (n = 6) were compared with a one-way ANOVA with a Holm-Bonferroni post-hoc correction. **(H)** V241R (V28R) completely disrupts ALX4 cooperativity. ALX4 and ALX4 V241R (V28R) protein (0, 37.5, 75, 150, and 300 nM) was combined with P3 fluorescent probe. Bars, dots, and error bars were defined in G. Tau factors (n = 12) were compared with an unpaired, two-sided student t-test. **(I)** ALX4 activates transcription in a cooperativity dependent manner. Luciferase assays were performed in transfected HEK293T cells. Bars denote average luciferase fold change of sample compared to reporter alone and dots represent an independent transfected well. Error bars denote standard deviation and luciferase fold changes between proteins were compared via a one-way ANOVA with a Tukey post-hoc correction. A single replicate of each EMSA is shown here and all replicates are shown in Supplementary Figure 6.

We first used PDBePISA to map the interactions of each ALX4 protein to DNA ^31^; (Figure 2E). Consistent with other HD proteins, two regions of the ALX4 HD make most of the base-specific interactions with DNA: the third helix (residues 254-269) contacts the major groove, and the N-terminal ARM motif (residues 214-218) contacts the minor groove (Figure 2D). Of these interactions, the major groove DNA contacts are largely symmetrical between the two ALX4 chains (Figure 2E; Supplementary Figure 5C-D). N264 (canonical HD numbering: N51) forms specific hydrogen bonds with TA**<u>A</u>**T (dA6/dA23) (Chain A DNA contact/Chain B DNA contact), and Y238 (Y25), R244 (R31), R257 (R44), R266 (R53), and R270 (R57) form identical non-specific contacts to the DNA backbone of their respective half-site (Figure 2E; Supplementary Figure 5C-D). However, the ALX4 Q263 (Q50) residue has distinct chain-specific configurations that contribute to major groove DNA binding. Prior studies have shown that the 50^th^ residue of the HD can influence the specificity of the TAAT**<u>NN</u>** nucleotides ^10,11,32,33^, and in the ALX4 dimer structure, Chain B: Q263 (Q50) has the potential to form specific contacts with dC9, dA10, dT24, and dT25 in the major groove. In contrast, Q263 (Q50) of Chain A does not form DNA contacts and instead forms intrachain hydrogen bonds with Q259 (Q46) (Supplementary Figure 5E-G), albeit at this resolution we cannot formally exclude the possibility that Q263 (Q50) of Chain A forms water mediated specific contacts with DNA.

As opposed to the largely symmetrical interactions found in the second and third helices, the ALX4 N-terminal ARM mediates several unique interactions apart from R218 (R5), which makes similar essential minor groove DNA contacts to each DNA half site. In the N-terminal ARM of Chain A, R215 (R2) inserts into the minor groove of DNA and forms a specific contact with dA6 and dT30 (Figure 2E-F), while R216 (R3) of Chain A forms a salt bridge with E255 (E42) of Chain B (described below). Hence, in this interface, R215 (R2) forms the key specificity creating contacts to DNA, whereas R216 (R3) forms critical protein-protein interactions. In comparison, the R215 (R2) residue in Chain B is positioned outside of the minor groove, and the side chain makes no DNA contacts (Figure 2E, G). This residue primarily faces the protein-to-protein interface and forms contacts with the main chains of Y238 (Y25) and V241 (V28) (described below). In its stead, the R216 (R3) side chain of Chain B inserts into the minor groove and forms specific contacts with dT13 and dA23 as well as a non-specific contact with dA15 (Figure 2E, G). Interestingly, there was not sufficient electron density to model the residues N210 (N −4) through G212 (G −3) and the side chains of K213 (K −1) and K214 (K1), suggesting that these regions were disordered in Chain B. It is possible that the R215 (R2) interaction to DNA stabilizes these N-terminal residues in Chain A. Thus, unlike the similar DNA-protein interactions mediated by the second and third helices, the N-terminal ARM of the two ALX4 proteins uses distinct residues to bind the minor groove of the P3 binding site.

### ALX4 dimer interactions are mediated via three interfaces

The two ALX4 proteins bound to DNA interact through three interfaces (Figure 3). The first interface is between the N-terminal ARM of Chain A and the turn between helix 1 and helix 2 on Chain B. In this interface, the K213 (K −1) side chain of Chain A forms hydrogen bonds with T236 (T23) and Y238 (Y25) of Chain B; the main chain of K214 (K1) of Chain A forms a hydrogen bond with V241 (V28) of Chain B; and R216 (R3) of Chain A forms a salt bridge with E255 (E42) of Chain B (Figure 3A-B), while R215 (R2) faces away from the interface and inserts into the DNA minor groove (Figure 2F). The second interface features head-to-head interactions between the start of the third helices where A256 (A43) form symmetrical van der Waals interactions as well as the salt bridge between R216 (R3) of Chain A and E255 (E42) of Chain B (Figure 3C-D). The third interface consists of the turn between helix 1 and helix 2 of Chain A and the N-terminal ARM of Chain B. Unlike in interface 1, R216 (R3) is inserted into the minor groove and forms no polar contacts with Chain A, whereas K213 (K −1) and K214 (K1) lack sufficient electron density to model their side chains. Hence, these three residues have little involvement in the protein-to-protein interface. Instead, R215 (R2) of Chain B forms hydrogen bonds with Y238 (Y25) and V241 (V28) of Chain A. In addition, N217 (N4) of Chain B forms a hydrogen bond with E245 (E32) of Chain A (Figure 3E-F). Thus, while interfaces 1 and 3 use symmetrical regions of the ALX4 protein, the residues utilized to facilitate these interactions are vastly different.

### R215 and R216 are critical for cooperative DNA binding to P3 sites

Given their chain-specific protein-protein and protein-DNA contributions, we next wanted to evaluate the roles that R215 (R2) and R216 (R3) have in cooperativity and DNA binding. We designed independent R215A (R2A) and R216A (R3A) mutations as well as a R215A/R216A (R2A/R3A) double mutant in the ALX4 DNA binding domain and evaluated their ability to cooperatively bind a P3 DNA sequence. Note, each ALX4 mutation tested in this manuscript was found to not significantly decrease protein stability as compared to wild type ALX4 in thermal stability assays unless specifically noted (Table 1). Through quantitative EMSAs, we found that R215A (R2A) and R216A (R3A) reduced cooperativity 5-fold and 10-fold, respectively (Figure 3G). Moreover, the effects of the N-terminal ARM mutations compound as the double mutant: R215A/R216A (R2A/R3A) reduced cooperativity 20-fold (Figure 3G).

**Table 1.** Melting temperatures derived from differential scanning fluorimetry for ALX4 structural mutations and ALX4 disease variants. Proteins were tested in triplicate, and melting temperatures are reported as mean ± standard deviation. Melting temperatures were compared via a one-way ANOVA with a Tukey post-hoc correction. The p-value of the variant compared to the wildtype ALX4 protein is shown. The melting curves are shown in Supplementary Figure 11A.

| Protein | Melting temperature |  |
| --- | --- | --- |
| | $^{\circ}\text{C}$ | *p-value* |
| ALX4 | 58.4±0.05 | - |
| R215A | 59.4±0.12 | 0.013 |
| R216A | 59.5±0.16 | 6.18e-3 |
| R215A; R216A | 59.2±0.39 | 0.065 |
| V241R | 61.4±0.08 | 3.38e-10 |
| K211E | 58.8±0.12 | 0.806 |
| R216G | 59.2±0.05 | 0.045 |
| R218Q | 58.9±0.05 | 0.576 |
| Q225E | 49.3±0 | 2.33e-14 |
| R272P | 56.8±0.21 | 1.54e- 5 |

Next, we assessed the role of the ALX4 R215 (R2) and R216 (R3) residues in DNA binding affinity using isothermal titration calorimetry (ITC) to a probe with a monomer binding site. Intriguingly, we found that the R215A (R2A) and R216A (R3A) mutations that significantly decreased cooperative DNA binding (Figure 3G) did not significantly impact ALX4’s ability to bind DNA (Table 2; Supplementary Figure 10). However, the double mutant decreased DNA binding affinity ∼14-fold (Table 2; p = 4.76e-10). The lack of affinity changes in the R215A (R2A) and R216A (R3A) mutations supports the idea that either arginine residue can contribute to monomeric DNA binding by making similar minor groove contacts (Figure 2G-H). However, the simultaneous mutation of both positions results in a loss of the A**<u>T</u>**TA minor groove DNA contacts, leading to a substantial loss in DNA binding affinity.

**Table 2.**
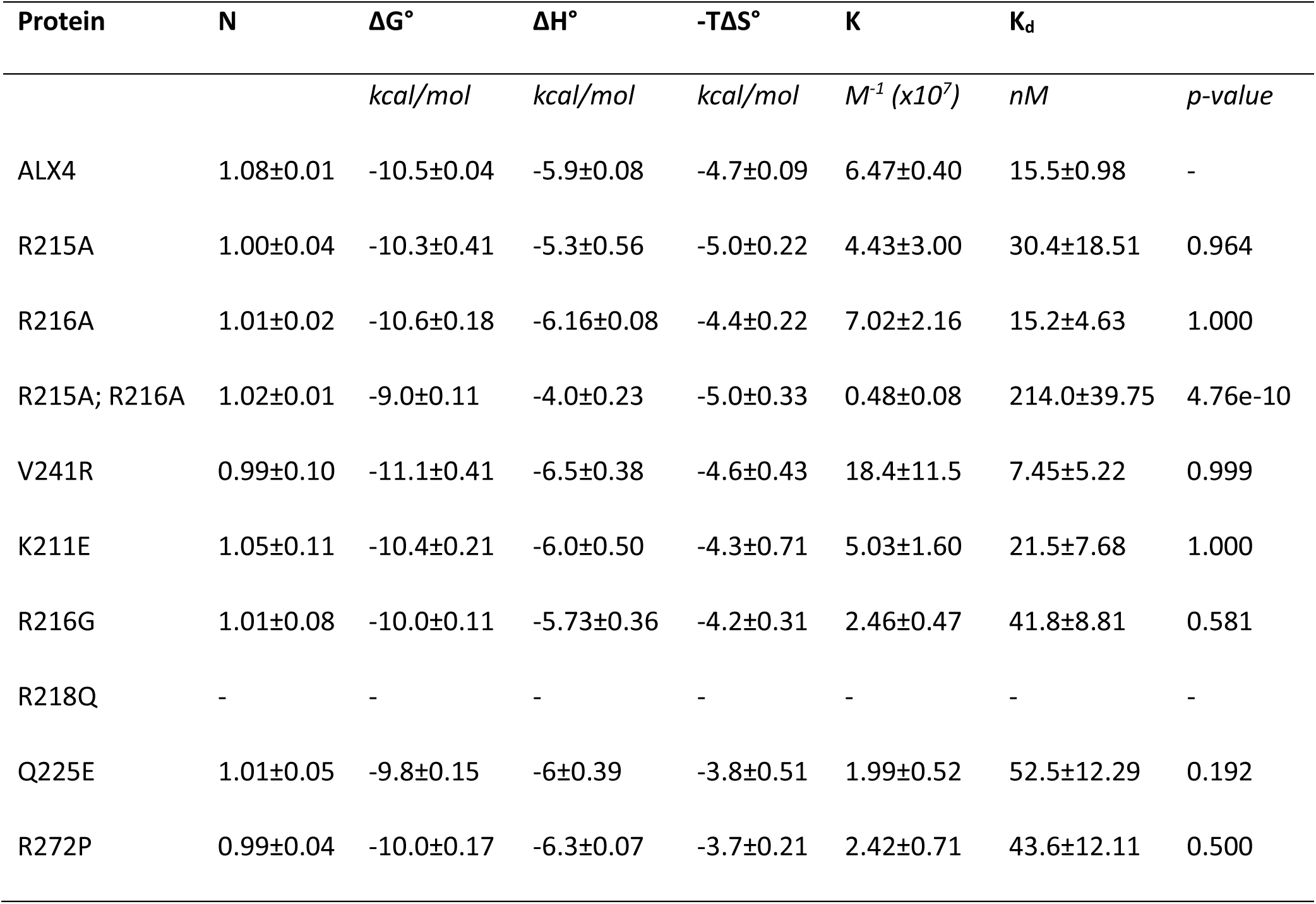
Calorimetric binding data of ALX4 structural mutations and ALX4 disease variants on the 14mer oligo duplex: CGCTAATTAGCTCG. All proteins were tested in triplicate, and measurements are reported as mean ± standard deviation. Affinities (K_d_) were compared via a one-way ANOVA with a Tukey post-hoc correction. The p-value of the variant compared to the wildtype ALX4 protein is shown. The thermograms are shown in Supplementary Figure 10. (N: stoichiometry; ΔG°: Gibb’s free energy; ΔH°: enthalpy; T: temperature (K); ΔS°: entropy; K: association constant; K_d_ : dissociation constant).

To further test the role of key protein-protein interactions in cooperative DNA binding, we next mutated the V241 (V28) residue that forms key hydrogen bonds and nonpolar interactions with either the R215 (R2) or R216 (R3) residues of the N-terminal ARM of the opposite chain. Using a V241R (V28R) mutation that inserts a larger charged residue predicted to sterically clash with the N-terminal ARM and disrupt these inter-protein interactions, we found that V241R (V28R) completely disrupted cooperativity and even hindered the ability of the second protein to independently bind to the second site as reflected by the Tau value being less than 1 (Tau = 0.327 ± 0.385) (Figure 3H). These data are consistent with prior studies showing that an I28R mutation disrupts cooperativity of the Prd S50Q protein to a P3 site ^15^. Importantly, the V241R (V28R) mutation did not significantly decrease monomer DNA binding affinity in ITC assays (Table 2), highlighting that this mutation selectively disrupts cooperativity.

### ALX4 mediated transcriptional activation via P3 sites is cooperativity dependent

Prior studies showed that ALX4 activates synthetic target sites containing 3 iterative - P3 sites in a spacer-dependent manner in luciferase reporter assays ^11,18^. To determine if transcriptional activation was cooperativity dependent rather than only DNA site dependent, we multimerized and cloned either 3 high affinity P3 sites (3xP3) or P4 sites (3xP4) in front of a minimal promoter and luciferase gene (schematic shown Figure 3I) and tested the ability of different ALX4 mutations to activate gene expression in HEK293T cells. As expected, wild type ALX4 activated the 3xP3 reporter but failed to activate the non-cooperative 3xP4 reporter (Figure 3I), demonstrating dimer site-dependent activation. Moreover, we found that the cooperativity-deficient ALX4 V241R (V28R) protein failed to activate either the 3xP3 or 3xP4 reporter (Figure 3I). Importantly, comparative Western blot and immunostaining assays revealed similar protein levels and nuclear localization of the wild type and V241R (V28R) ALX4 proteins in cell culture (Supplementary Figure 6F). Taken together, these data demonstrate that ALX4 transcriptional activation is cooperativity dependent.

### The ALX4 dimer structure combines interactions found in the PRD and Engrailed structures

To better understand how the protein-protein and protein-DNA interactions in the ALX4 structure compare with other HD/DNA complexes, we analyzed the Prd-S50Q/DNA dimer structure on a similar P3 site ^15^, the Engrailed HD/DNA monomer structure ^5^, and the Aristaless (al) HD/DNA monomer structure ^34^ (Supplementary Figure 7). Overall, the major groove DNA interactions of these structures were very similar. However, the N-terminal ARM interactions were distinct between them due in large part to which Arg residue is inserted into the minor groove of DNA. Both the En and al monomer structures use the R3 residue to facilitate minor groove binding (Supplemental Figure 7), whereas the Paired S50Q protein uses the R2 residue to mediate highly similar DNA interactions thereby freeing the R3 residue to mediate largely symmetrical protein-protein interactions between the two Chains (Figure 4B, D). Intriguingly, the mechanisms used by ALX4, especially by the ALX4 N-terminal ARM, are a blend of both the Prd dimeric structure and the Engrailed and *al* monomeric structures. The protein-DNA interface at Chain A resembles the interface found in the Paired dimeric structure, in which R216 (R3) interacts with E255 (E42) rather than inserting into the minor groove. In this case, R215 (R2) compensates for R216 (R3) to make the necessary protein-DNA contacts. The protein-DNA interface at Chain B resembles the Engrailed and al monomeric structures, where R216 (R3) inserts into the minor groove and mediates the necessary contacts with DNA while R215 (R2) makes negligible DNA contacts. In this interface, the protein-protein interface is unique to ALX4 as R2 and N4 make no protein-protein contacts and minimal van der waals interactions in the Prd S50Q structure. Overall, these data highlight how the flexible use of which arginine residue in the N-terminal ARM is used to contact DNA can allow for other N-terminal ARM residues to contribute to cooperative DNA binding through either symmetric (Paired S50Q) or asymmetric (ALX4) interactions.

**Figure 4.**
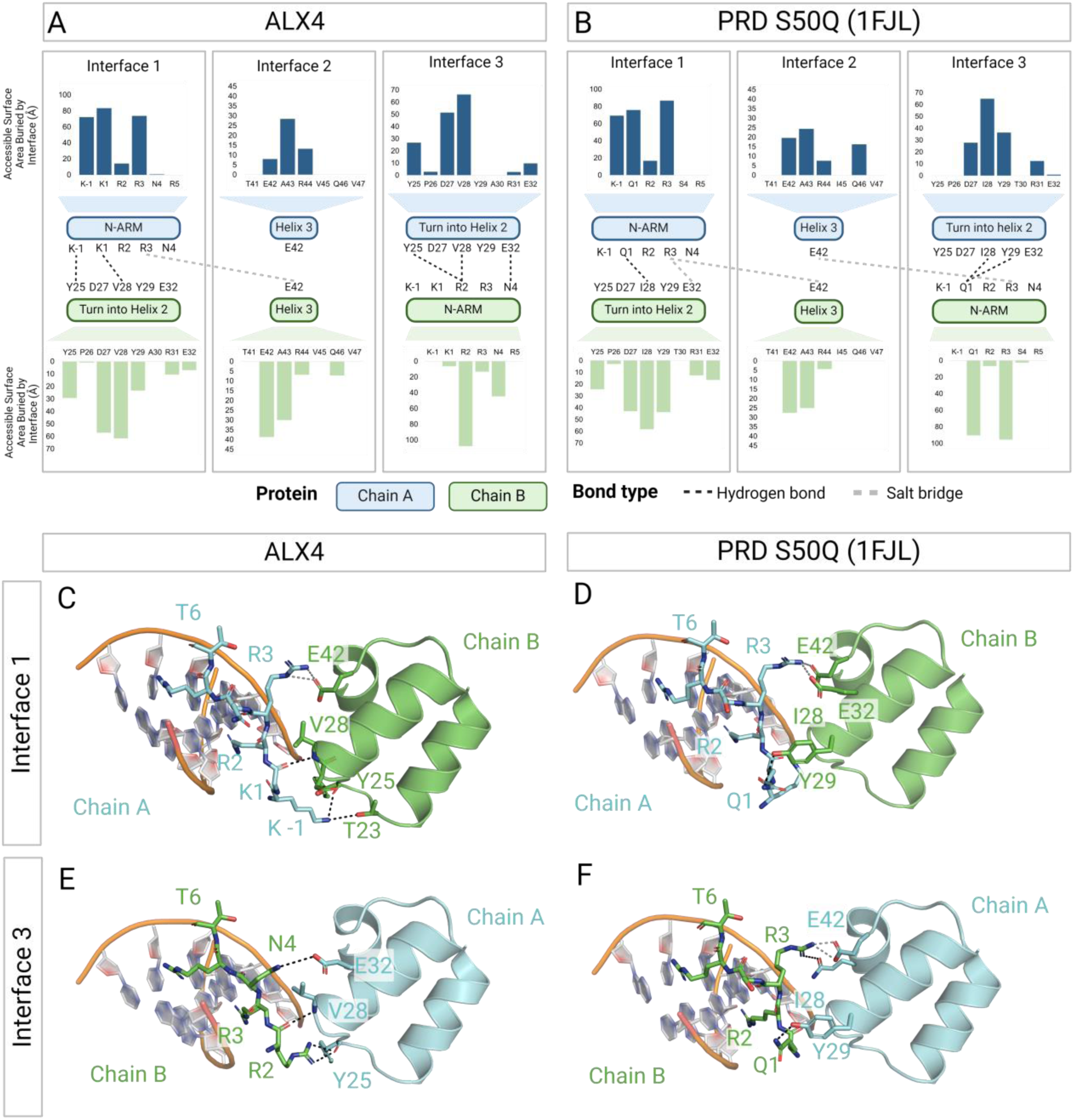
Comparison of ALX4 and PRD S50Q structures reveals asymmetric N-terminal ARM interactions in ALX4 but not PRD. **(A-B)** Protein-protein interactions of the ALX4 and PRD S50Q structure with the canonical HD numbering for comparison. Van der waals interactions are shown in the bar graphs and are measured by the residue’s accessible surface area buried by the other chain. Polar bond contacts are shown as dashed lines between contacting residues with hydrogen bonds in black and salt bridges in gray. **(A, C)** In Chain A, K −1 forms bonds with T23 and Y25, K1 contacts V28, and R3 contacts E42. R2 inserts into the minor groove. **(B, D)** This interface is quite similar to the PRD S50Q structure as Q1 interfaces with I28 and Y29 in the turn between helix 1 and 2, and R3 interfaces with E32 and E42 in helix 2 and 3. **(A, E)** K −1, K1 and R3 of Chain B have limited involvement in the protein-protein interface as R3 inserts into the minor groove. Instead, R2 contacts the main chains of Y25 and V28 in the turn between helix 1 and 2, and N4 interfaces with E32 in the second helix. **(B, F)** This interface is not consistent with the PRD structure as Q1 and R3 are again the primary facilitators of the protein-protein interactions between the N-terminal ARM and the turn between helix 1 and 2 as well as the beginning of helix 3.

### ALX4 variants associated with disease differentially impact cooperativity and DNA binding

With our improved understanding of the residues implicated in DNA binding and cooperativity, we next sought to classify the functional impacts of ALX4 missense variants associated with disease. There are three conditions associated with ALX4 missense variants: 1) Enlarged parietal foramina (PFM), 2) Sagittal craniosynostosis (CS), and 3) Frontonasal dysplasia (FND). Despite the large range of phenotypes and severities, limited functional characterization has been performed. Here, we assessed how ALX4 disease variants impact ALX4 function by measuring cooperativity with quantitative EMSAs (Supplementary Figure 8), DNA binding with ITC assays (Table 2; Supplementary Figure 10), protein stability with differential scanning fluorimetry assays (Table 1; Supplementary Figure 11A), and transcriptional activity with luciferase assays in transfected HEK293T cells. The expression of each ALX4 protein in HEK293T cells was confirmed via western blot (Supplementary Figure 8G), and protein translocation into the nucleus was evaluated via immunofluorescence (Supplementary Figure 9). In total, we characterized five ALX4 missense variants: four PFM variants (R216G, R218Q, Q225E, and R272P), one CS variant (K211E), and two FND variants (R216G and Q225E) (Figure 5A). Note, some of these variants have been reported to cause more than one disease.

**Figure 5.**
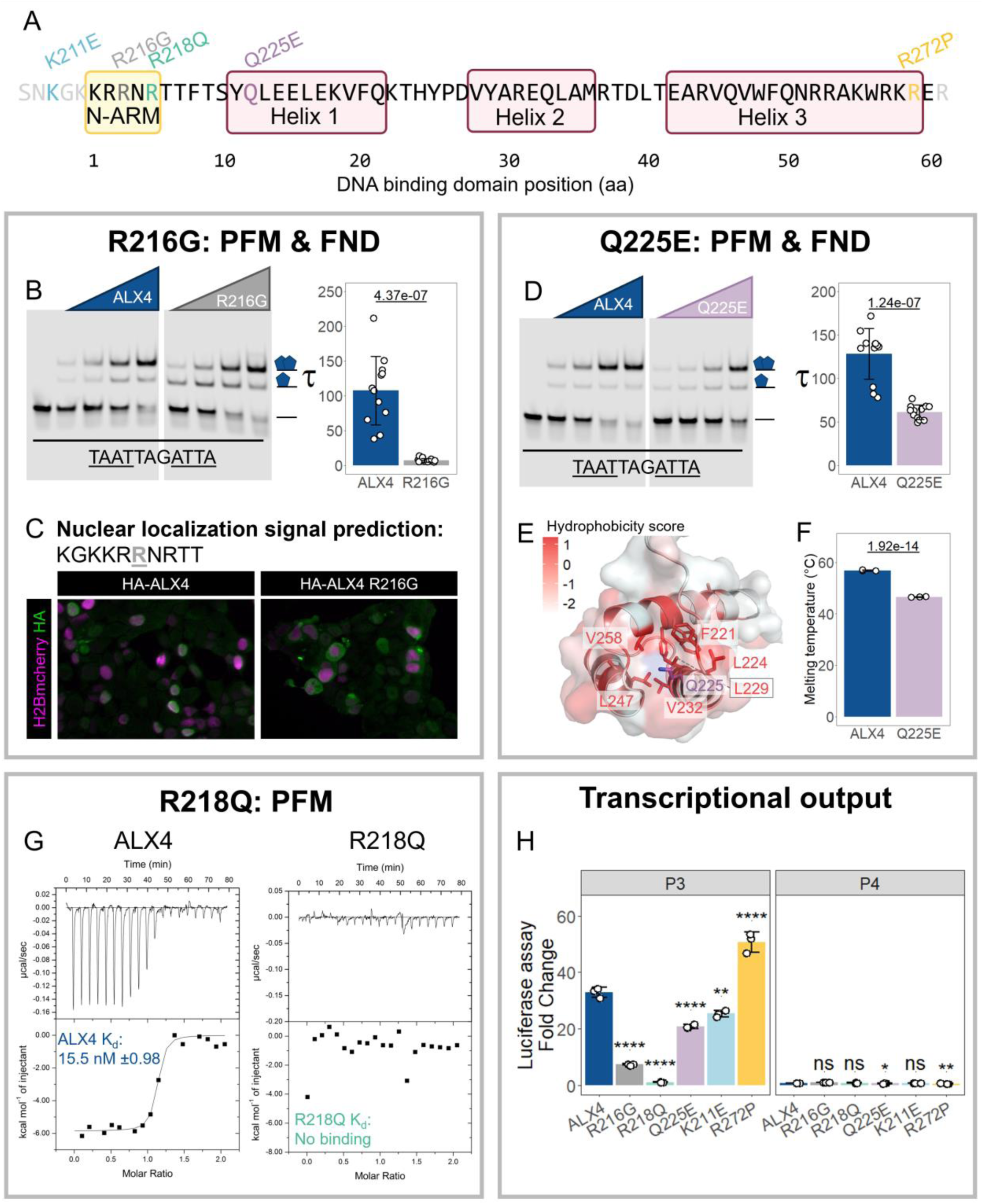
ALX4 human disease variants impact DNA binding and/or cooperativity which affect transcriptional output on a P3 site. **(A)** Five missense variants located within or near the DNA binding domain were characterized. **(B)** ALX4 R216G (R3G) disrupts cooperativity 10-fold. ALX4 and the corresponding variant protein (0, 37.5, 75, 150, and 300 nM) was combined with P3 fluorescent probe. A single replicate is shown but all replicates are shown in Supplementary Figure 8. Tau cooperativity factors were calculated for every lane in which protein was added. Bars depict the average Tau for each protein and each dot represents a Tau from an independent binding reaction (n = 12). Error bars denote standard deviation. Tau factors were compared with an unpaired, two-sided student t-test. **(C)** ALX4 R216G (R3G) hindered protein’s ability to localize in the nucleus. H2B-mcherry was used as a transfection control. NLS Mapper ^35^ predicted the only ALX4 nuclear localization signal to be the N-terminal ARM. **(D)** ALX4 Q225E (Q12E) reduced P3 mediated cooperativity 2-fold. See section B for Tau calculation details. **(E)** The Q225E (Q12E) variant is located near the hydrophobic core of ALX4. Protein schematic was created in PYMOL and residues are colored based on hydrophobicity. **(F)** ALX4 Q225E reduced protein stability. Melting temperatures derived from differential scanning fluorimetry were compared via a one-way ANOVA with a Tukey post-hoc correction. The p-value of the variant compared to the wildtype ALX4 protein is shown. The melting curves are shown in Supplementary Figure 11A. **(G)** ALX4 R218Q (R5Q) failed to bind the 14mer oligo duplex, CGCTAATTAGCTCG, in ITC whereas the wildtype ALX4 protein bound the sequence with an affinity of 15.5 nM. **(H)** Select ALX4 disease variants reduce transcriptional activation on the P3 site while no variant or wildtype activated the P4 site. Luciferase assays were performed in transfected HEK293T cells. Bars denote average luciferase fold change sample compared to reporter alone, while each dot represents an independent transfected well. Error bars denote standard deviation and luciferase fold changes between proteins were compared via a one-way ANOVA with a Tukey post-hoc correction where *p < 0.05, **p < 0.01, ***p < 0.001, ****p < 0.0001.

The ALX4 R216G (R3G) allele has been associated with both enlarged PFM and FND ^25^. Here, we found that R216G (R3G) reduces cooperativity 10-fold in EMSAs (Figure 5B) and this variant has no impact on DNA binding affinity in ITC assays (Table 2), much like the R216A (R3A) variant tested above (Figure 3G). The loss of cooperativity can be explained by the ALX4 crystal structure as R216 (R3) forms salt bridges with E255 (E42) on Interface 1 (Figure 3A). Further, the preservation of DNA binding is consistent with R215’s (R2) ability to insert into the minor groove of DNA to form the necessary base pair contacts and compensate for the loss of R216 (R3) (Figure 2F). Consistent with the importance of cooperativity in ALX4-mediated transcriptional activation, we found that the R216G variant resulted in a 10-fold reduction in luciferase output on the 3xP3 reporter (Figure 5H). Intriguingly, however, immunostaining of the ALX4 R216G variant revealed decreased nuclear localization in comparison to wild type ALX4, with the ALX4 R216G protein being observed in the cytoplasm and nucleus (Figure 5C). NLSMapper ^35^ predicted that the N-terminal ARM is the sole nuclear localization signal in ALX4. These data are consistent with previous studies of other HD proteins, including close homologs, CART1 and ARX, that found that the basic residues in the N-terminal ARM can function as a nuclear localization signal ^36–41^ Hence, the ALX4 R216G variant results in two molecular defects: decreased nuclear localization and decreased cooperative DNA binding to the P3 dimer site, both of which likely contribute to the reduction of ALX4-mediated activation on the P3 site.

The ALX4 Q225E (Q12E) variant is associated with PFM and FND ^21^. We found that this variant reduced cooperativity ∼2-fold (Figure 5D) with little change in DNA binding affinity (Table 2). This alteration in cooperativity modestly reduced transcriptional output on the P3 site (Figure 5H). A recent protein binding microarray found that the ALX4 Q225E variant weakens DNA binding but preferentially maintains binding to “AATAAA” 6mers ^42^. The authors suspected that this residue change disrupts protein stability or influences the hydrophobic core due to Q225E’s proximity to residues within the core (Figure 5E). In support of this idea, we found that the Q225E variant decreased protein stability in differential scanning fluorimetry compared to the wildtype protein (Figure 5F), while maintaining proper secondary structure content as measured by circular dichroism (Supplementary Figure 11B). However, the question of how this destabilization leads to a biased retention of specific kmers has not yet been addressed.

The ALX4 R218Q (R5Q) allele is a well-studied variant associated with PFM ^19,22,23^. We found that this variant abolishes all DNA binding in ITC assays (Figure 5G) as well as in EMSAs (Supplementary Figure 7B). Consistent with these results, the ALX4 R218Q variant failed to activate the 3xP3 reporter (Figure 5H), even though it was properly localized to the nucleus (Supplementary Figure 8). The complete loss of DNA binding is predicted by the structure as R218 forms critical bonds with the minor groove of DNA. Further, R5 in the DNA binding domain is conserved across most HD TFs ^10^ and is considered a disease variant hotspot for HDs ^43^. Consistent with these findings, the R218Q (R5Q) variant correlates with more severe defects than other ALX4 point mutations or deletions ^23^.

We also tested the K211E (K −3E) variant that has been associated with increased penetrance of CS ^26^ and the R272P (R59P) variant that has been associated with PFM ^24^. We found no substantial reductions in cooperativity, DNA binding affinity, protein stability or transcriptional output for these variants (Figure 5H; Table 1; Table 2). Interestingly, we did not recapitulate previous findings that K211E and R272P increased and decreased transcriptional output, respectively. However, this discrepancy may be cell-type specific as we used HEK293T cells, whereas past studies used calvarial osteoblasts ^26^. It is also important to note that these variants occur on surface accessible residues on the protein. Thus, while these residues did not mediate protein-to-protein interactions in the ALX4 dimer structure, they could be implicated in protein-protein interactions with other co-factors.

## Discussion

In this work, we defined the mechanisms underlying how the Paired-like factor, ALX4, binds cooperatively to the TAAT – NNN – ATTA (P3) dimer site and assessed how known disease variants impact DNA binding and transcriptional regulation. Using available genomic binding data, we first found that ALX4 binds to several distinct motifs and of these only ALX4 binding to the P3 dimer site is independent of the craniofacial master regulator, TWIST1. This contrasts with ALX4’s ability to bind the heterodimeric coordinator motif and independent monomer sites, both of which were dependent on TWIST1 occupancy and/or TWIST1 induced accessibility. Second, we used crystallography to solve the ALX4 structure on a P3 site, providing new insights into the protein-DNA and protein-protein interactions that mediate ALX4 cooperativity and DNA binding specificity. Third, we characterized the functional impacts of five ALX4 disease variants associated with PFM, FND, or increased penetrance of CS using a combination of quantitative DNA binding and transcriptional reporter assays. Collectively, these studies provide considerable new insights into the molecular mechanisms underlying HD DNA binding specificity and the diverse molecular defects caused by disease-associated ALX4 alleles.

### The diverse and flexible roles of the N-terminal ARM in mediating cooperativity and DNA binding specificity

The way in which HDs gain sufficient DNA binding specificity to regulate their distinct *in vivo* targets while using highly conserved DNA binding domains with highly similar *in vitro* DNA binding preferences has been a long-standing question in the field. Cooperativity addresses one part of this paradox as dimerization increases TF-DNA binding specificity in three ways. (i) Cooperativity typically involves binding longer DNA binding sites (i.e. the 11 bp P3 dimer site) that are statistically less likely to occur at random compared to a single 4 to 6 bp monomer site. (ii) A TF that cooperatively binds to a specific dimer site is likely to better compete with other TFs that bind the independent monomer sites. For example, we previously found that the HD, Gsx2, cooperatively bound a TAAT – 7N – TAAT dimer site with higher affinity compared to its monomer site ^44^. This added affinity would provide Gsx2 a selective advantage in binding this site over non-cooperative HDs, whereas ALX4 would have a similar advantage on the P3 dimer site. (iii) Since cooperativity typically involves protein-protein contacts, the number of residues that contribute to TF-DNA binding specificity increases. In Supplementary Figure 7, we highlight the residues implicated in DNA side chain contacts for the monomeric Engrailed and aristaless structures. Unsurprisingly, the key residues: R3, R5, and N51 are highly conserved across all *Drosophila* ^10^ and human HDs ^42^, consistent with most HDs binding the same monomer sites in *in vitro* assays ^2–4^. Here, we identified seven additional HD residue positions that indirectly contribute to ALX4 DNA specificity by facilitating P3 site cooperativity. These residues: K1, R2, N4, T23, V28, E32, and E42, are largely conserved across the Paired-like subclass ^45^, explaining why other Paired-like factors demonstrate similar cooperative DNA binding behaviors ^11,14,46,47^. However, these residues have little conservation across the entire HD family, highlighting why only select HDs can bind to the P3 site cooperatively ^14^. Interestingly, AlphaFold3 ^48^ correctly predicted that ALX4 would bind to the TAAT DNA binding sites within the P3 site, however AlphaFold3 docked the two proteins in a head-to-tail orientation rather than the correct head-to-head orientation (Supplementary Figure 12). Because of this, AlphaFold would be unable to correctly predict the interfacing regions and residues, highlighting both the utility in solving the crystal structures of these complexes and the current limitations of using a prediction program to determine protein-protein interfaces.

An unexpected finding from the ALX4 dimer structure was the asymmetric use of the N-terminal ARM residues to mediate DNA-protein and protein-protein interactions. Given the palindromic nature of the P3 site (TAAT −3N – ATTA), one would have predicted that ALX4 would use similar mechanisms to bind each half site, much like was previously found for the Prd S50Q dimeric structure ^15^. However, we found that the N-terminal ARM residues facilitating these interfaces are unique between the two structures. In the Prd S50Q structure, interfaces 1 and 3 mediate protein-protein interactions that are largely symmetrical and are facilitated by positions 1 and 3 in the N-terminal ARM (Figure 4) ^15^. In contrast, interface 3 in the ALX4 structure is driven by R2, N4, Y25, and E32, revealing a newfound importance for these residues in P3 site cooperativity (Figure 4E). R2, N4, and Y25 do not make protein-protein contacts in the Prd S50Q structure and were not previously thought to be essential for cooperativity. Further, R2, Y25, and E32 are conserved across the Paired-like class ^45^, whereas N4 is not well conserved. ^49,50^ Future studies will be required to address the importance of the fourth position in contributing to P3-mediated cooperativity.

Comparative studies across HDs reveal that there is some flexibility as to which arginine residue in the N-terminal ARM mediates direct minor groove DNA binding contacts. The N-terminal ARM consists of several arginine residues that contribute to DNA binding specificity. While all HDs use the highly conserved R5 residue to make specific contacts, additional contacts are typically mediated by a second arginine residue at either position 2 or position 3. We found that the asymmetry between the two ALX4 proteins is largely driven by whether R2 or R3 inserts into the minor groove. Moreover, we showed that while both arginine residues are required for cooperativity (Figure 3G), either arginine is sufficient for high affinity monomer DNA binding (Table 2). The preservation of DNA binding after R2A or R3A mutations suggests that either R2 or R3 can insert into the minor groove and compensate for the loss of the other, but by doing so, the sole arginine can no longer facilitate the protein-protein interactions required to maintain cooperativity. R2 and R3’s ability to make similar DNA contacts has been shown previously: A bacterial one-hybrid survey of 84 *Drosophila* HDs revealed that both positions have the potential to create specificity contacts with T**<u>A</u>**AT ^32^. This has been structurally confirmed as well as R3 makes specificity contacts in Engrailed ^5^, PITX2 ^51^, and aristaless ^34^, whereas Msx-1 uses R2 ^52^. Interestingly, R2 and R3 are conserved throughout all the Paired-like class ^45^, but many other HDs have a lysine in one of these positions ^10,42^. While lysine has a similar charge as arginine, lysine residues insert into DNA minor grooves ∼3-fold less than arginine residues ^53^ and appear less frequently in protein-protein interfaces than arginine residues ^54^. Thus, these structural studies on ALX4 and other HD proteins highlight how differences in key HD residues, especially within the N-terminal ARM, contribute to distinct DNA binding and cooperativity behaviors.

### ALX4-induced transcriptional activation is dependent on TF cooperativity

ALX4 variants, including several missense variants within the HD, have been associated with craniofacial birth defects such as PFM, FND, and increased penetrance of CS. How such mutations disrupt the ability of ALX4 to bind DNA, regulate target gene expression, and ultimately cause disease is not well understood. Here, we used the insight gained from our ALX4 dimer structure to interrogate the ability of several HD missense alleles to bind cooperative dimer DNA binding sites as well as monomer sites. Combining quantitative DNA binding assays with a dimer-dependent transcriptional assay, we were able to identify molecular defects for three of five tested ALX4 HD missense variants. Based on their locations in the ALX4 crystal structure, it was not surprising that neither K211E nor R272P significantly impacted monomer or cooperative dimer binding. Moreover, both proteins were properly localized and activated transcription similarly as wild type ALX4, making it unclear how these variants cause disease. In contrast, the other three disease variants have clear molecular defects that impact DNA binding. R218Q (R5Q) completely abolished monomer and dimer DNA binding, whereas the R216G (R3G) and Q225E (Q12E) disease variants selectively decreased cooperative DNA binding. In addition, the R216G and Q225E variants caused secondary deficits that are likely to contribute to their observed ∼10-fold and ∼2-fold reduced transcriptional output, respectively (Figure 5). R216G reduced protein nuclear localization (Figure 5C) and Q225E impacted protein stability (Figure 5E-F). Thus, these three ALX4 HD disease variants have distinct molecular defects that impact their ability to activate transcription via cooperative P3 binding sites.

Past work showed that ALX4 only transcriptionally activates reporter genes containing the P3 site, and not monomer, P2, P4, or P5 sites ^18,19^. Consistent with these data, we found that a cooperativity-deficient ALX4 protein, ALX4 V241R (V28R), could no longer transcriptionally activate the P3 site, despite maintaining its ability to bind DNA (Figure 3H-I). The correlation between cooperativity and transcriptional output emphasizes the importance of cooperativity on ALX4 function and suggests that ALX4 requires a partner to enact a transcriptional change. A recent study found that ALX4 cooperatively binds with TWIST1 to regulate human craniofacial development ^28^, providing a second case in which ALX4 binds with a partner to induce a transcriptional change. In alignment with this idea, we found that ALX4 binding to the coordinator, HD monomer, and E-box site were dependent on TWIST1 (Figure 1), whereas ALX4 binding to the P3 site was not dependent on TWIST1 binding (Figure 1; Figure 3I) ^18,19^. This behavior extends to close homologue, CART1 (sometimes referred to as ALX1) as it too specifically activates transcription on a P3 site ^18^, and has high co-occupancy with TWIST1 in differentiated hCNCCs ^28^. Other HDs have also been shown to alter transcriptional activity via homo-dimeric cooperativity. For example, the ability of CRX, another Paired-like HD protein, to mediate transcriptional changes in massively paralleled reporter assays significantly correlated with P3 dimer sites, but not monomer sites ^13^. In addition, the Gsx2 HD protein was found to induce opposing transcriptional outcomes on dimer (gene stimulation) versus monomer (repression) sites ^44^. Thus, cooperativity-dependent transcriptional activation is not unique to ALX4, but a common mechanism used by many HDs.

## 4. Methods

### Genomic analyses

The ALX4 and TWIST1 genomic binding data was acquired from Kim et al., 2024 and the accession IDs for these data are listed in the Supplementary table 2. Sequencing data underwent adapter trimming with Cutadapt (v4.4) ^55^ and quality control with fastqc (v0.11.2) ^56^ via the wrapper trimgalore (v0.6.6) ^57^. Sequences were mapped to hg19 with bowtie2 (v2.3.4.1) with the following settings: --local --very-sensitive-local --no-unal --no-mixed --no-discordant ^58^. Results were deduplicated with picard MarkDuplicates ^59^. MACS3 with the --keep-dup all --qvalue 0.01 options was used for peak calling of all genomic assays ^60^. For C&R peak calling, --call-summits --SPMR settings were also applied; for H3K27ac ChIP-seq peak calling, --broad --broad-cutoff 0.1 settings were also applied, and for ATAC-seq peak calling --call-summits --SPMR --shift -75 --extsize 150 settings were also applied. BigWigs and heatmaps were generated using deepTools ^61^. C&R and ChIP-seq bigwigs were normalized with the log2 ratio of the immunoprecipitated datasets versus the IgG controls. ATAC-seq bigwigs were normalized to reads per genomic content.

Homer *de novo* motif analysis identified 6, 12, and 18bp length enriched sequences in the ALX4 C&R datasets ^30^. annotatePeaks.pl from the Homer package was used to annotate the coordinator, dimer, monomer, and E-box sites within the ALX4 bound regions. For footprint analysis, genome coverage tool from bedtools calculated 5’ read coverage of reads that were less than 120bp in length ^62^. We determined whether a predicted site was bound by comparing the MNase digestion signal of the motif (which should be protected when bound by a TF) to regions flanking the motif (which should be highly accessible) within a set motif window. The set motif window for the coordinator, dimer, monomer, and E-box site was −10bp to 10bp, −6bp to 6bp, −3bp to 3bp, and −3bp to 3bp respectively, where 0 is the motif center. The motif flank was ±40bp outside of the motif window. The following equation was used to determine if a site was bound, where *M*_*f*_stands for the average 5’ binding signal of the 40bp flanking either side of the motif window and *M*_*w*_ stands for the average 5’ binding signal of the motif window: 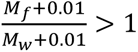 ^44^. The 0.01 addition eliminated any errors introduced by dividing by zero. We used the GRanges package in R to remove any overlapping or palindromic repeat sites as well as any single sites that were part of a bound composite site ^63^. Supplementary Figure 3A shows the number of sites that were removed in each of the described steps.

To statistically compare the reads per million of the ALX4 C&R in wildtype conditions versus the ALX4 C&R in TWIST1 depleted conditions in Figure 1E, ALX4 C&R in TWIST1 depleted conditions was normalized to the ALX4 C&R in wildtype conditions by multiplying the reads per million by a scaling factor derived by the *E.coli* spike-in control ^28^.

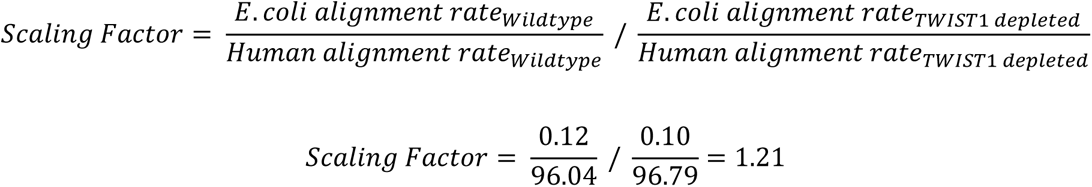

All custom code can be found at https://github.com/cainbn97/ALX4_footprinting_analysis.

### Protein preparation

The mouse ALX4 sub-fragment was PCR amplified using Accuzyme DNA polymerase (Bioline) from mALX4 cDNA (Genscript: OMu19476) and cloned into pET14P. Note: while mouse cDNAs were used, the mouse and human ALX4 protein sequences are identical for the regions used in this study. pET14P is a modified pET14b plasmid in which a PreScission Protease site was inserted between the 6xHis-tag and TF coding sequence ^14^. PCR primers used for amplification included a C-terminal Strep-tag and are listed in Supplementary Information. ALX4 variant cDNAs were prepared via site directed mutagenesis. All pET14P-ALX4 constructs were transformed into C41(DE3) (Sigma-Aldrich) *E. coli*, grown in LB media, and induced with 0.1mM IPTG. Following bacterial cell lysis, ALX4 proteins were purified using a combination of Ni-NTA and Strep-Tactin affinity, and size exclusion chromatography. The N-terminal 6xHis-tag was removed via digestion with the PreScission Protease (GE Healthcare) as previously described ^14^. Protein purity was confirmed via SDS-PAGE with GelCode blue staining (Thermo Scientific) (Supplementary Figure 6C; Supplementary Figure 8F), and protein concentrations were assessed by absorbance at UV 280nm.

### Isothermal Titration Calorimetry

ITC experiments were performed using a Microcal VP-ITC microcalorimeter. For all experiments, the DNA duplexes were placed in the syringe at ∼100µM, and all ALX4 proteins were placed in the cell at ∼10µM. Titrations consisted of an initial 1µl injection followed by nineteen 14µl injections. All experiments were performed in a buffer containing 50mM sodium phosphate, pH 6.5 and 150mM NaCl at 20^◦^C. All samples were dialyzed overnight to ensure buffer match. c values (c=KMn) for all ITC experiments were 200<c<702. All binding experiments were performed in triplicate. Final raw data were analyzed using ORIGIN and fit to a one-site binding model.

### Crystallography

ALX4-DNA complexes were formed prior to crystallization by mixing purified protein in a 2:1 (protein:DNA) ratio with a final complex concentration of ∼10mg/ml (∼1mM). The DNA used for crystallization was a 17-mer duplex, containing the sequence 5’ – CGC**<u>TAAT</u>**TCA**<u>ATTA</u>**ACG – 3’ with two ALX4 binding sites (bold, underlined) spaced three base pairs apart. The ALX4-DNA complex crystalized in a solution containing 0.1M Tris pH 7.5, 0.2M Trimethylamine N-oxide, and 20% PEG MME 2000. The subsequent crystals diffracted to 2.39 Å resolution and belong to the space group P3_2_21 with cell dimensions a=b=70.46 c=159.14 Å. The asymmetric unit of the crystal contained two copies of ALX4 and one copy of the DNA duplex. Unbound ALX4 was crystallized at 10mg/ml in a solution containing 0.1M MgCl_2_, 0.1M KCl, 0.1M PIPES pH=7.0, 20% PEG Smear Medium. The crystals diffracted to 2.39 Å resolution and belong to the space group I2_1_3 with cell dimensions a=b=c=95.78 Å. All synchrotron diffraction data was collected at APS LS-CAT.

### Structure Determination, Model Building, and Refinement

Phaser was used for molecular replacement with the PDB file 1FJL as the search model ^64^. COOT was used for manual model building within the observed electron density ^65^ and Phenix was used for refinement with TLS parameters ^66^. The final ALX4/DNA and ALX4 models were refined to a R_work_/ R_free_= 22%/24% and 21%/25%, respectively, with good overall geometry and validated with MolProbity ^67^. Protein structure schematics were created by PYMOL v2.5.7 ^68^. Accessible surface area buried by interface measurements and polar interactions were calculated by PDBePISA v1.52 ^31^.

### Circular Dichroism

CD experiments were performed on an Aviv Circular Dichroism Spectrophotometer 215 using a 0.5 mm quartz cuvette (Hellma Analytics). The cuvette was not removed during a series of scans taken from 300 to 190 nm in 1 nm increments. Proteins were dialyzed into a buffer containing 5 mM sodium phosphate (pH 6.5) and 150 mM NaF, and then diluted to the desired concentration of ∼0.30 mg/ml in the same buffer. Data are plotted as mean residue ellipticity, θ, in units of deg cm2 /dmol per residue.

### Electrophoretic mobility shift assays

EMSA probes were prepared by annealing a 5’-IRDye 700nM labeled oligo (IDT) to an oligo containing the binding sites of interest as previously described ^69^. Sequences used for probe preparation are listed in the supplementary information. EMSA binding reactions were prepared as previously described with 34 nM of denoted fluorescent probe ^70^. Binding reactions were incubated at room temperature for 20 minutes. 4% polyacrylamide gels were run at 150V for 2 hours. Gels were imaged on the Li-Cor Odyssey CLx scanner and band intensity was measured via the Li-Cor image studio software. Tau was calculated by the below equation, where *[P_2_D]* represents dimer binding, *[PD]* represents monomer binding, and *[D]* represents unbound probe as previously described ^12^. This equation was derived by the equilibrium reactions of a single protein binding DNA as well as the equilibrium equation of the binding of a second protein to the monomeric complex.

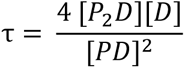

### Luciferase assays

Full-length human ALX4 cDNA sequences of the wildtype and variants were cloned into a HA-tagged pcDNA3.1 mammalian expression vector by Genscript. cDNA sequences are listed in the supplementary information. 5 UAS sites, 3 P3 or P4 sites, and the minimal promoter from the *Drosophila* Ac gene were cloned between KpnI and NcoI sites of pGL3-basic (Promega). Note, sites were cloned 3’ from 5 upstream activation sequence (UAS) sites. However, Gal4 was not added, thus these sites had no impact on luciferase output. Reporter sequences are provided in supplementary information.

Forty thousand HEK293T cells were cultured in a 48-well plate in DMEM and 20% fetal bovine serum for 18 hours prior to transfection. Each well was transfected with 80 ng of total DNA: 5 ng of Renilla plasmid, 25 ng of indicated firefly reporter plasmid, and the denoted amount of ALX4 cDNA and pcDNA3.1 empty expression construct using the Effectene Transfection reagent (Qiagen) following manufacturer’s protocol. Cells were lysed with 80 µL of passive lysis buffer (Promega) 30 hours after transfection. Firefly luciferase and Renilla luciferase florescent values were measured with the Promega Dual Luciferase Assay kit (Promega: E1910) and the GloMax-96 Microplate Luminometer. Firefly luciferase was normalized to Renilla luciferase to control for transfection efficiency. All results are reported as expression fold change over the reporter alone sample. All samples were run in triplicate.

### Western blot

Four hundred thousand HEK293T cells were cultured in a 6-well plate in DMEM and 20% fetal bovine serum for 18 hours prior to transfection. Each well was transfected with 600 ng of ALX4 cDNA in pcDNA3.1 or a pcDNA3.1 empty expression construct with Effectene Transfection reagent (Qiagen) following the manufacturer’s protocol. Cells were lysed by mechanically scraping cells in 750 µL of radioimmunoprecipitation assay (RIPA) buffer (150 mM of NaCl, 50 mM of Tris HCl pH 8.0, 1% NP-40, 0.5% sodium deoxycholate, 0.1% sodium dodecyl sulfate, 1x cOmplete Protease Inhibitor Cocktail (Roche: 04693116001), and 1 mM of phenylmethylsulfonyl fluoride). Lysates were incubated on ice for 5 minutes and then sonicated for 5 seconds with 25 seconds of rest time for a total of 3 sonication cycles.

Lysates were run on a 4-20% gradient acrylamide gel (BioRad) at 100V for 90 minutes. Gels were transferred to a polyvinylidene difluoride (PVDF) membrane via a semi-dry transfer that was run for 1 hour at 15V at room temperature. Membranes were blocked in a 0.5% Casein solution for 1 hour at room temperature. Membranes were probed with HA-tag (Roche: 3F10; 1:250) and β-actin (Li-Cor 926-42212; 1:2000) overnight at 4°C in PBS + 0.05% Tween-20 (PBST) + 0.05% Casein. The membrane was then washed 3 times with PBST and then incubated with 1:5000 IRDye 680RD Goat anti-Rat (Li-Cor: 926-68076) and IRDye 800CW Donkey anti-Mouse (Li-Cor: 926-32212) in PBST + 0.05% Casein for 1.5 hours at room temperature. The membrane was then washed 3 times in PBST and imaged on the Li-Cor Odyssey CLx scanner.

### Cell immunofluorescence

Forty thousand HEK293T cells were cultured in a 48-well plate in DMEM and 20% fetal bovine serum for 18 hours prior to transfection. Each well was transfected with 40 ng of an H2B-mcherry expression vector to label transfected cells and 40 ng of the indicated ALX4 cDNA in pcDNA3.1. Thirty hours after transfection, cells were washed with PBS and then fixed by incubating cells in 4% paraformaldehyde for 10 minutes at 4C. Cells were permeabilized in PBS + 0.5% Triton-X for 5 minutes at room temperature. Cells were blocked in PBS + 5% normal donkey serum (Jackson ImmunoResearch Laboratories Inc.: 017-000-121) for 1 hour at room temperature and then probed with the HA-tag (Roche: 3F10; 1:4000) in PBS + 0.2% Tween-20 + 5% normal donkey serum overnight at 4°C. Cells were washed three times in PBS + 0.2% Tween-20 and then incubated in Alexa Fluor 488 Donkey Anti-Rat (Invitrogen: A-21208; 1:1000) in PBS + 0.2% Tween-20 + 5% normal donkey serum for 1 hour at room temperature. Cells were then washed three times in PBS + 0.2% Tween-20 in which DAPI (1:1000) was added to the second wash. Cells were imaged on a Nikon ECLIPSE Ts2R Inverted Microscope. The LUT of the H2Bmcherry was changed from yellow to magenta in FIJI to allow for the discrimination of channels in the merged images.

## 6. Data availability

The structures have been deposited into the Protein Data Bank with accession numbers 9D9R and 9D9V.

## 7. Acknowledgements

Figures were created with Biorender.com.

## 8. Author contributions

This scientific study was conceived and planned by B.C., R.A.K, and B.G. Genomic analyses were performed by B.C. All construct synthesis and protein purification were carried out by Z.Y. All crystallography and X-ray structural studies were conducted by Z.Y. and analyzed by Z.Y. and R.A.K. Protein structure visuals and schematics were generated by B.C. ITC, differential scanning fluorimetry assays, and CD assays were performed by Z.Y. EMSAs were performed by B.C. and E.T. Cell immunofluorescence assays, western blots, and reporter assays were performed by B.C. The manuscript was written by B.C. and edited by R.A.K. and B.G.

