## Supplemental Information for "Structural and biochemical characterization of the ALX4 dimer reveals novel insights into how disease alleles impact ALX4 function"

### 1. Supplementary figures

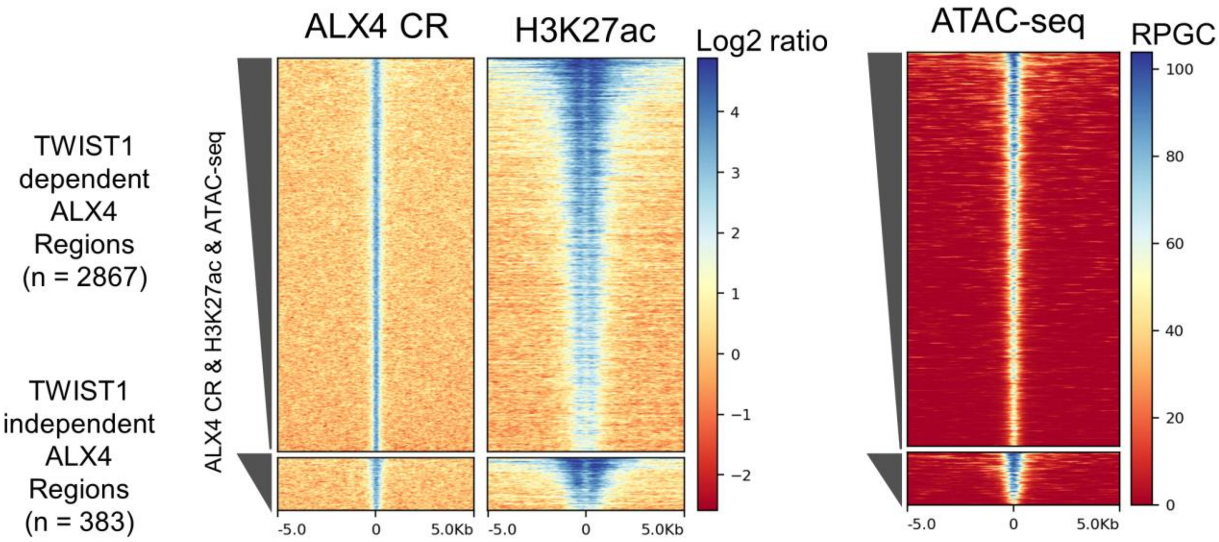


Supplementary Figure 1. The 3250 ALX4 bound regions in hCNCCs are acetylated and accessible. These regions were categorized into TWIST1 dependent and TWIST1 independent regions by comparing the ALX4 binding signal in a TWIST1+ versus TWIST1- background (Figure 1B). ALX4 CR and H3K27ac ChIP signals were normalized to their IgG controls by a log2 ratio. ATAC-seq signal was normalized to reads per effective genomic content (RPGC). Regions were sorted by ALX4 CR, H3K27ac, and ATAC-seq signal.

### 6 bp *de novo*

| Rank | Motif | P-value | log10 p-value | % of Targets | % of Background | |
|---|---|---|---|---|---|---|
| 1 | 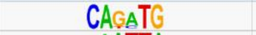 | 1.00e-359 | -8.27E+02 | 63.82% | 30.94% | E-box |
| 2 | 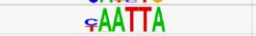 | 1.00E-249 | -5.74E+02 | 56.45% | 29.38% | Monomer |
| 3 | 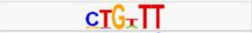 | 1.00E-84 | -1.93E+02 | 67.48% | 51.46% | |
| 4 | 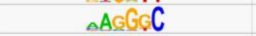 | 1.00E-43 | -1.00E+02 | 59.38% | 47.84% | |
| 5 | 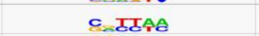 | 1.00E-40 | -9.43E+01 | 62.38% | 51.22% | |
| 6 | 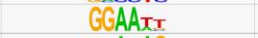 | 1.00E-36 | -8.41E+01 | 56.94% | 46.37% | |
| 7 | 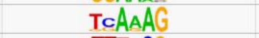 | 1.00E-28 | -6.65E+01 | 68.28% | 59.21% | |
| 8 | 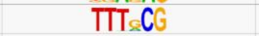 | 1.00E-18 | -4.35E+01 | 18.96% | 13.57% | |
| 9 | 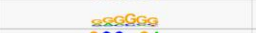 | 1.00E-17 | -4.04E+01 | 43.69% | 36.67% | |
| 10 | 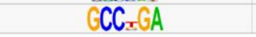 | 1.00E-09 | -2.17E+01 | 20.88% | 16.92% | |

### 12 bp *de novo*

| Rank | Motif | P-value | log10 p-value | % of Targets | % of Background | |
|---|---|---|---|---|---|---|
| 1 | 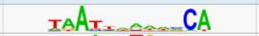 | 1.00e-818 | -1.88E+03 | 52.48% | 11.01% | |
| 2 | 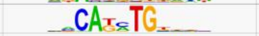 | 1.00e-556 | -1.28E+03 | 48.57% | 13.51% | |
| 3 | 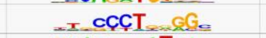 | 1.00E-81 | -1.88E+02 | 23.79% | 12.20% | |
| 4 | 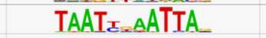 | 1.00E-76 | -1.76E+02 | 4.16% | 0.56% | Dimer |
| 5 |  | 1.00E-35 | -8.23E+01 | 0.50% | 0.00% |  |
| 6 | 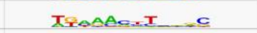 | 1.00E-34 | -8.05E+01 | 0.61% | 0.01% | |
| 7 | 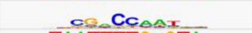 | 1.00E-34 | -7.95E+01 | 0.55% | 0.01% | |
| 8 | 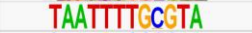 | 1.00E-33 | -7.68E+01 | 0.47% | 0.00% | |
| 9 | 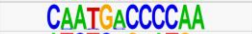 | 1.00E-33 | -7.68E+01 | 0.47% | 0.00% | |
| 10 | 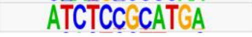 | 1.00E-33 | -7.68E+01 | 0.47% | 0.00% | |

### 18 bp *de novo*

| Rank | Motif | P-value | log10 p-value | % of Targets | % of Background | |
|---|---|---|---|---|---|---|
| 1 | 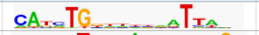 | 1.00e-2221 | -5.12E+03 | 59.63% | 3.06% | Coordinator |
| 2 | 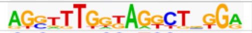 | 1.00E-45 | -1.05E+02 | 0.61% | 0.00% | |
| 3 | 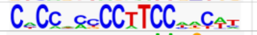 | 1.00E-42 | -9.89E+01 | 0.58% | 0.00% | |
| 4 | 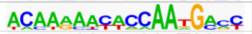 | 1.00E-42 | -9.89E+01 | 0.58% | 0.00% | |
| 5 | 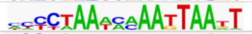 | 1.00E-40 | -9.33E+01 | 0.55% | 0.00% | |
| 6 | 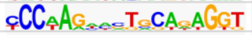 | 1.00E-40 | -9.33E+01 | 0.55% | 0.00% | |
| 7 | 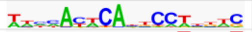 | 1.00E-38 | -8.97E+01 | 0.67% | 0.01% | |
| 8 | 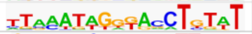 | 1.00E-38 | -8.78E+01 | 0.53% | 0.00% | |
| 9 | 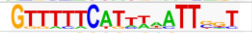 | 1.00E-36 | -8.44E+01 | 0.58% | 0.01% | |
| 10 | 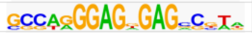 | 1.00E-35 | -8.23E+01 | 0.50% | 0.00% | |

Supplementary Figure 2. Homer *de novo* motif analysis of ALX4 CUT&RUN showed enrichment of four distinct site types.

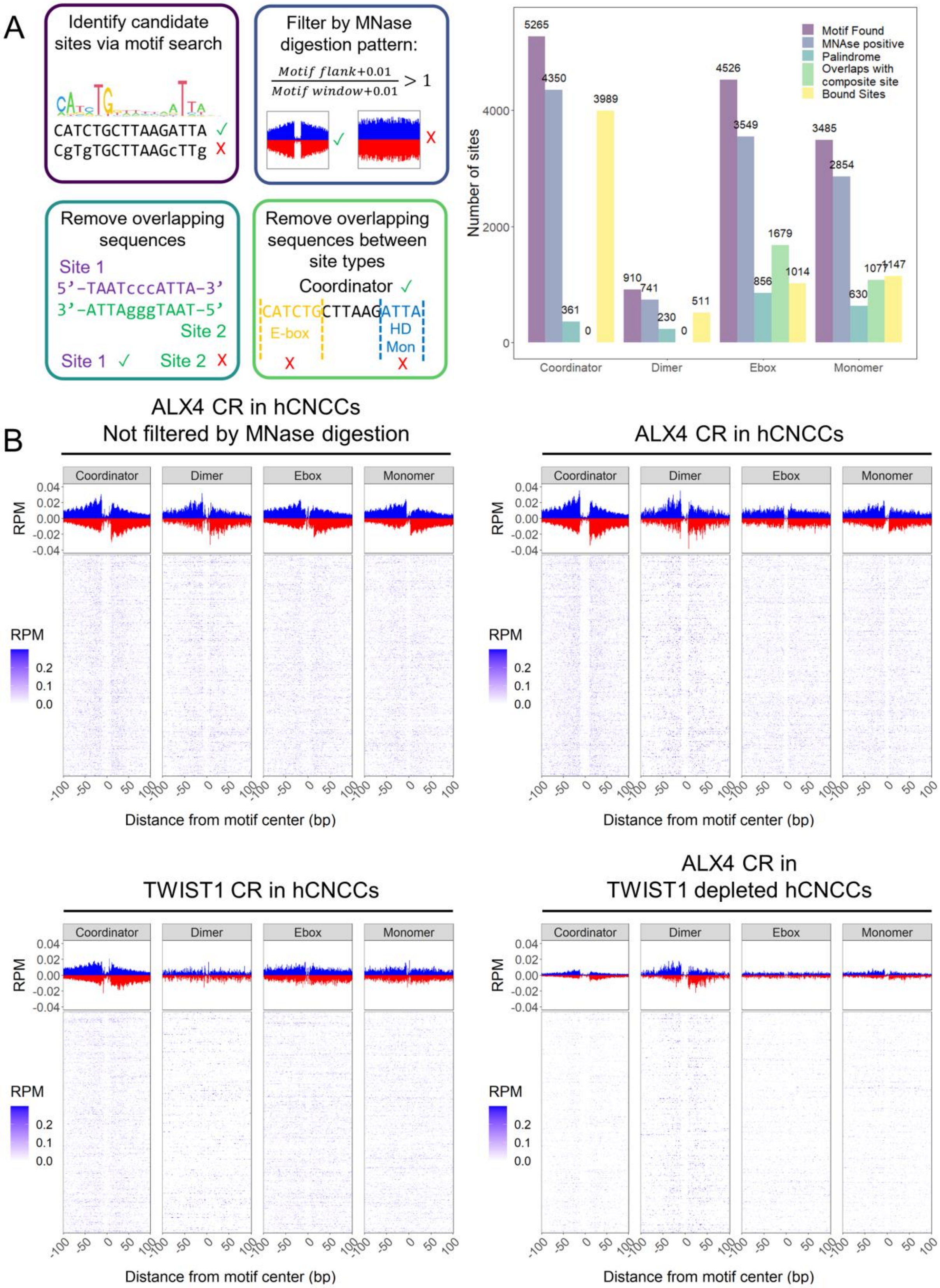
A
Identify candidate sites via motif search
CATCTGCTTAAGATTA ✓
CgTgTGCTTAAGcTTg X
Filter by MNase digestion pattern:
Motif flank+0.01 / Motif window+0.01 > 1
Remove overlapping sequences
Site 1
5'-TAATcccATTA-3'
3'-ATTAgggTAAT-5'
Site 2
Site 1 ✓ Site 2 X
Remove overlapping sequences between site types
Coordinator ✓
CATCTGCTTAAGATTA
E-box
HD Mon
Motif Found
MNAse positive
Palindrome
Overlaps with composite site
Bound Sites
Number of sites
5265
4350
361
0
3989
910
741
230
0
511
4526
3549
856
1679
1014
3485
2854
630
1077
1147
Coordinator
Dimer
Ebox
Monomer
B
ALX4 CR in hCNCCs
Not filtered by MNase digestion
ALX4 CR in hCNCCs
TWIST1 CR in hCNCCs
ALX4 CR in TWIST1 depleted hCNCCs
RPM
Distance from motif center (bp)

12

Supplementary Figure 3. Four site types from 3 CR assays were independently footprinted. **(A)** Sites were found via motif search of the 3250 ALX4 bound regions with Homer. Sites were then filtered by MNAse digestion pattern where the ratio of MNAse digestion signal flanking the motif (±40 bp region flanking the motif window was used) and MNAse digestion within the motif window (±10 bp for the coordinator, ±6 for the dimer, and ±3 for the E-box and monomer site) had to be greater than 1 for the site to be considered bound. Sites that contained duplicated palindromes were removed and any sites that were part of a larger composite site were removed. Note, most sites that were found to have a motif were bound. **(B)** Average digestion patterns (bar graph) and digestion patterns for each site (heatmap) are shown for the ALX4 CR footprint prior to MNase digestion filtering as well as the footprint after MNAse digestion filtering of the ALX4 CR, TWIST1 CR, and ALX4 CR in a TWIST1 depleted background datasets. Note, the bar graphs denote strand specific digestion whereas the heatmaps show the sum of both strands at a given position.

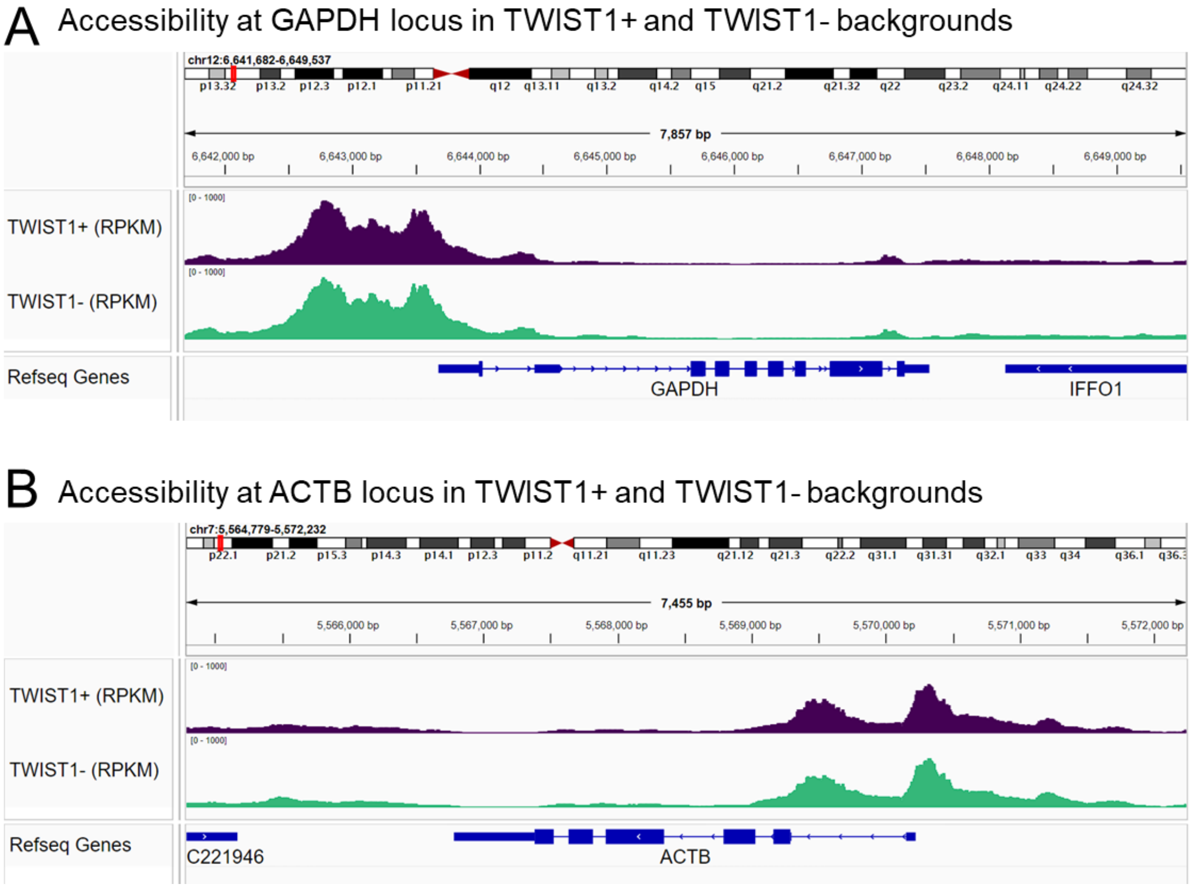


Supplementary Figure 4. TWIST1+ and TWIST- ATAC-seq datasets have similar signal to noise. Signal track views of housekeeping genes, **(A)** GAPDH and **(B)** ACTB, demonstrate similar signal to noise between TWIST1+ ATAC-seq and TWIST1- ATAC-seq datasets. Signal tracks were normalized by reads per kilobase per million mapped reads (RPKM) and were created with the Integrative Genomics Viewer [71].

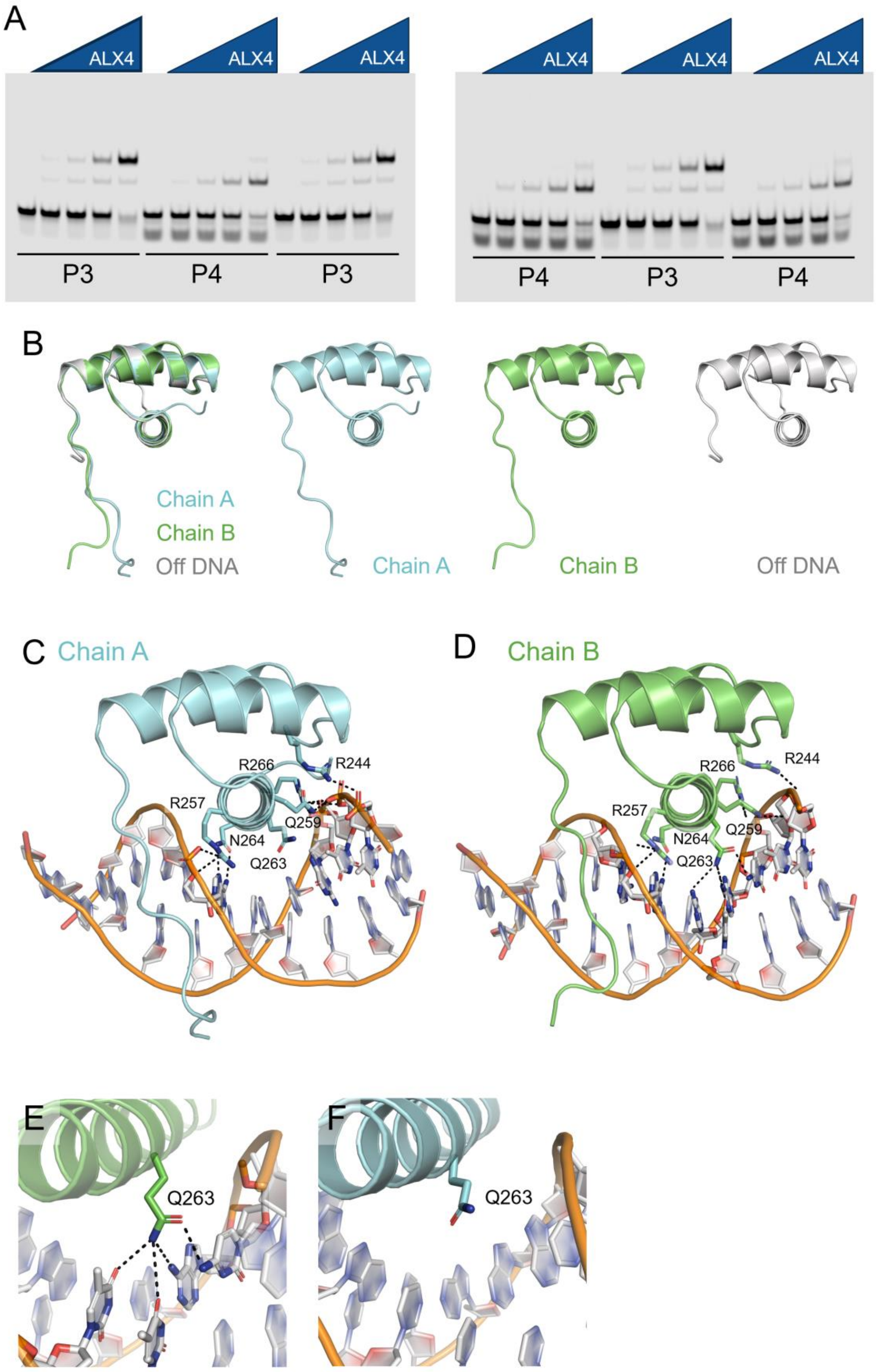
A
ALX4
ALX4
ALX4
ALX4
ALX4
ALX4
P3
P4
P3
P4
P3
P4
B
Chain A
Chain B
Off DNA
Chain A
Chain B
Off DNA
C
Chain A
R266
R244
R257
Q259
N264
Q263
D
Chain B
R266
R244
R257
N264
Q259
Q263
E
Q263
F
Q263

31

Supplementary Figure 5 ***(A).*** All replicates for quantitative EMSAs used for Tau measurements in Figure *2*B. All binding reactions contain 34 nM of respective fluorescent probe where the P3 probe contains the TAAT – TAG – ATTA sequence and the P4 probe contains the TAAT – TAGG – ATTA sequence. The ALX4 HD binds cooperatively in a spacer length dependent manner. ALX4 protein was tested in triplicate at 0, 37.5, 75, 150, and 300 nM concentrations. **(B)** There are no structural changes in the three alpha-helices between the two ALX4 HD proteins bound in the dimer complex and the ALX4 protein not bound to DNA. However, the N-terminal ARM differs in conformation between these three chains and is only partially resolved in the free ALX4 structure. **(C-D)** Schematics comparing protein-DNA interactions of **(C)** ALX4 Chain A and **(D)** ALX4 Chain B. N264 of both chains makes symmetrical specific contacts to TA**A**T, whereas R244 (R31), R257 (R44), R266 (R53), and R270 (R57) form identical non-specific contacts with the DNA backbone. **(E)** Q263 (Q50) on Chain B has the potential to form specificity contacts with dC9, dA10, dT24, and dT25 in the major groove. **(G)** However, Q263 (Q50) of Chain A forms no specific contacts.

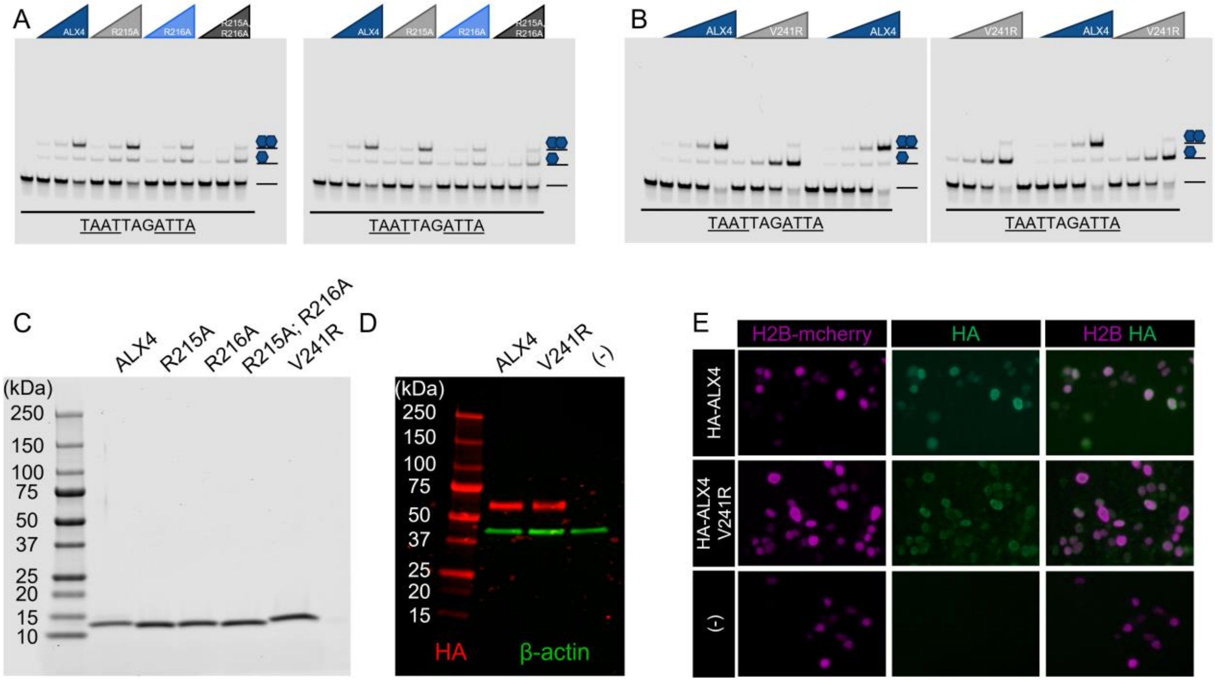


Supplementary Figure 6. **(A-B)** All replicates for quantitative EMSAs used for Tau measurements in Figure 3G-H. All binding reactions contain 34 nM of respective fluorescent probe. Schematics of the protein-DNA complexes are shown to the right of the gel. **(A)** N-terminal ARM mutations negatively impact ALX4 cooperative binding to the P3 site. ALX4 WT and structural mutant protein were tested in duplicate at 0, 50, 100, and 200 nM concentrations. **(B)** ALX4 V241R (V28R) fully disrupts cooperativity. ALX4 and ALX4 V241R (V28R) protein was tested in triplicate at 0, 37.5, 75, 150, and 300 nM concentrations. **(C)** Protein purity was assessed with a Coomassie stained SDS-PAGE. 5 µM of protein was loaded into each lane. **(D)** Western blot confirming protein expression in transfected HEK293T lysates. 600 ng of HA-tagged full-length ALX4 cDNA was transfected into a single well of a 6-well plate. β-actin was used as a loading control. HA (Roche: 3F10) staining is shown in red and β-actin (Licor: 926-42212) is shown in green. **(E)** Nuclear localization of ALX4 and ALX4 V241R in transfected HEK293T cells. 40 ng of H2B-mcherry and 40 ng of corresponding HA tagged ALX4 cDNA in pcDNA3.1 were transfected into HEK293T cells. H2B-mcherry was used as a transfection control and its autofluorescence is shown in magenta. HA (Roche: 3F10) staining is shown in green. Note: brightness and contrast were individually adjusted to best match H2B and HA expression levels to determine localization. Expression levels cannot be gleaned from these images and were instead measured via western blot in (F).

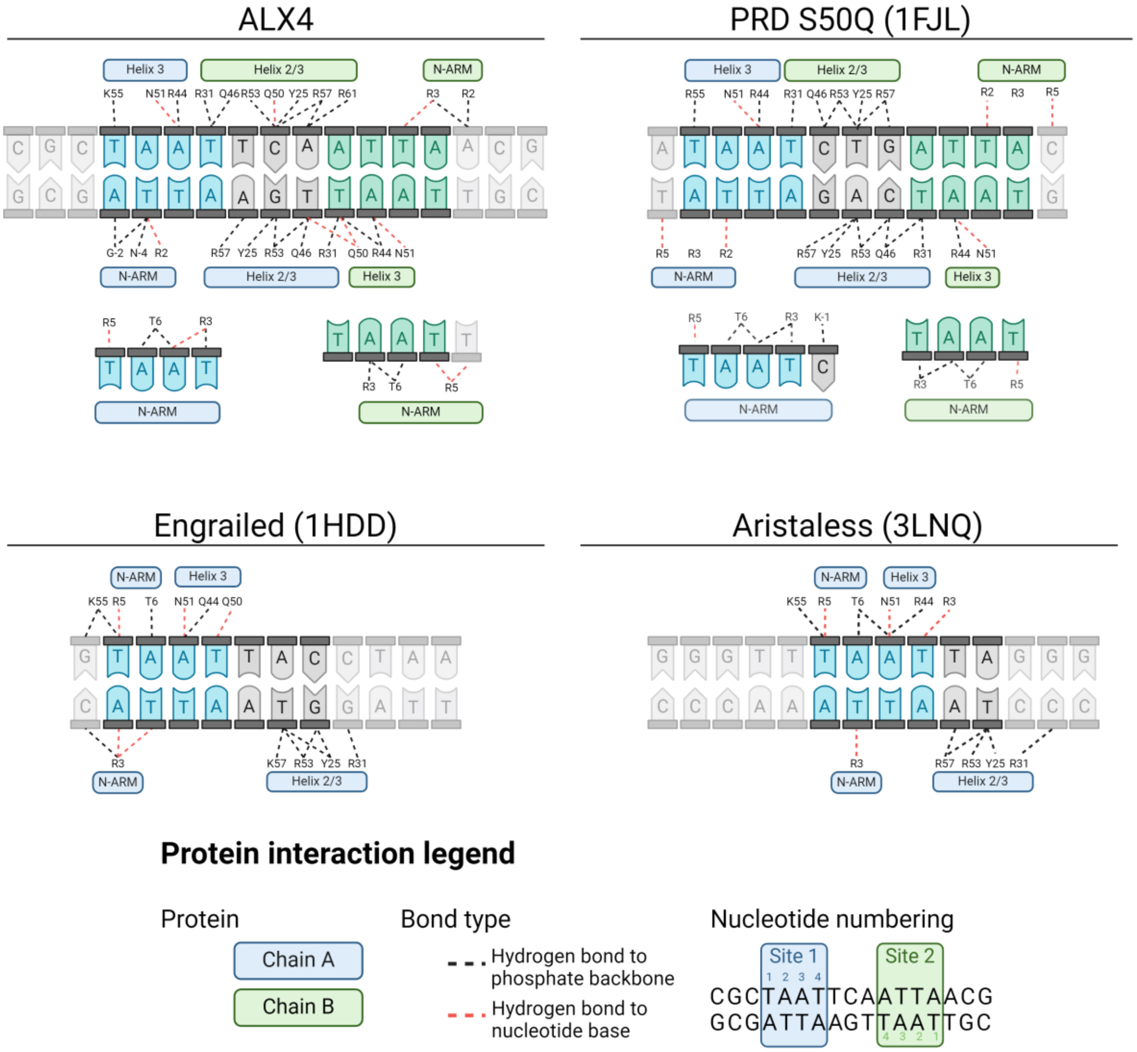


Supplementary Figure 7. Four HD-DNA structures exhibit similar helix 3-DNA interactions. However, the N-terminal ARM interactions are distinct between them. PISAePDB was used to determine polar contacts. For the ALX4 and PRD S50Q structures, Chain A interactions are shown in blue and Chain B interactions are shown in green. Polar contacts are shown as dashed lines between the protein residue and DNA base. Black lines indicate contacts to the backbone and red lines indicate contacts to the nucleotide base.

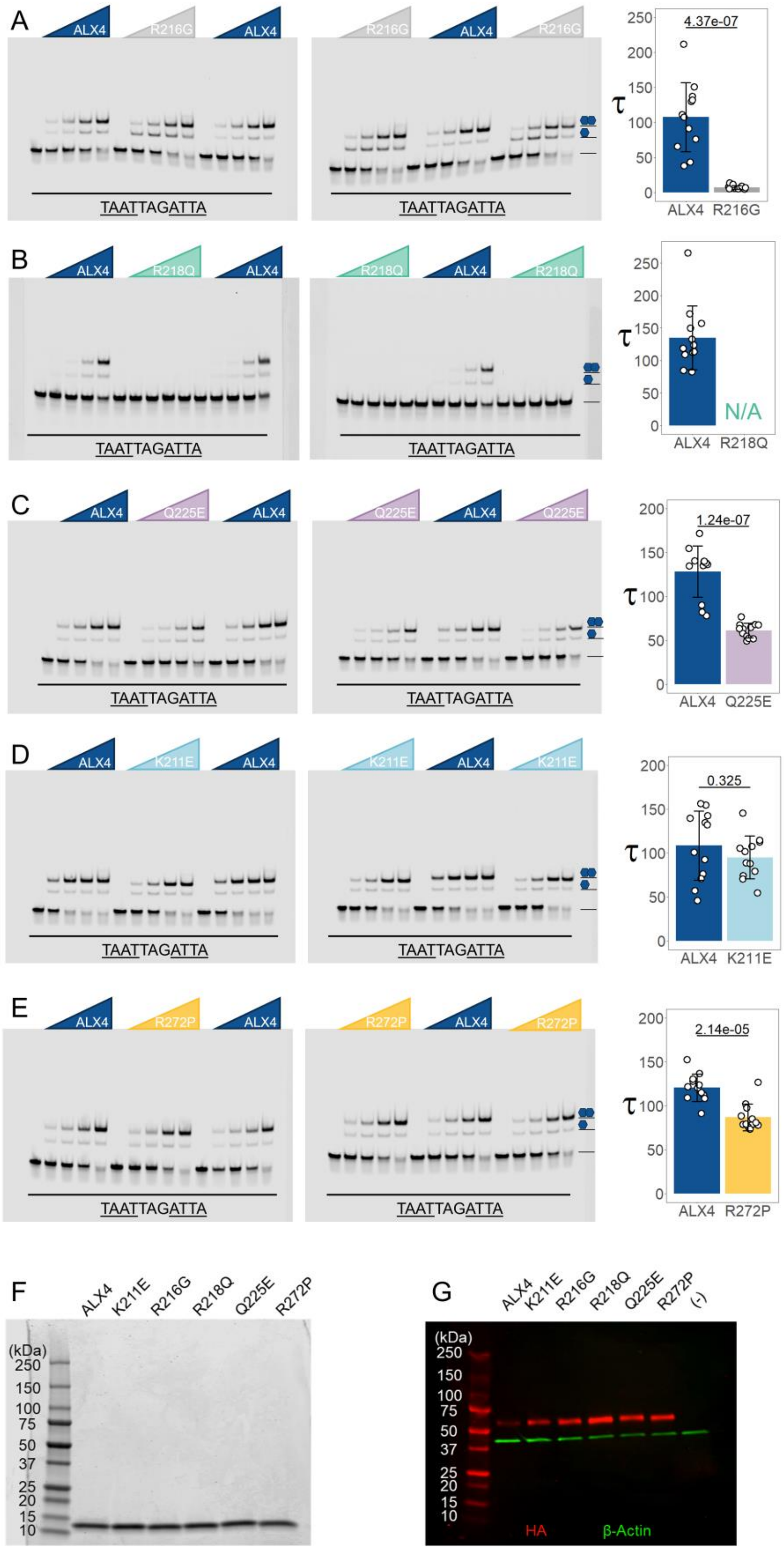
A
ALX4
R216G
ALX4
R216G
ALX4
R216G
TAATTAGATTA
TAATTAGATTA
4.37e-07
τ
250
200
150
100
50
0
ALX4
R216G
B
ALX4
R218Q
ALX4
R218Q
ALX4
R218Q
TAATTAGATTA
TAATTAGATTA
N/A
ALX4
R218Q
C
ALX4
Q225E
ALX4
Q225E
ALX4
Q225E
TAATTAGATTA
TAATTAGATTA
1.24e-07
ALX4
Q225E
D
ALX4
K211E
ALX4
K211E
ALX4
K211E
TAATTAGATTA
TAATTAGATTA
0.325
ALX4
K211E
E
ALX4
R272P
ALX4
R272P
ALX4
R272P
TAATTAGATTA
TAATTAGATTA
2.14e-05
ALX4
R272P
F
ALX4
K211E
R216G
R218Q
Q225E
R272P
(kDa)
250
150
100
75
50
37
25
20
15
10
G
ALX4
K211E
R216G
R218Q
Q225E
R272P
(-)
(kDa)
250
150
100
75
50
37
25
20
15
10
HA
β-Actin

Supplementary Figure 8. **(A-E)** All replicates for quantitative EMSAs used for Tau measurements in Figure 5. All binding reactions contain 34 nM of P3 probe. Schematics of the protein-DNA complexes are shown to the right of the gel. ALX4 and ALX4 disease variants were tested at 0, 37.5, 75, 150, and 300 nM concentrations. Tau cooperativity factors were calculated for every lane in which protein was added. Bars depict the average Tau for each probe and each dot represents a Tau from an independent binding reaction (n = 12). Error bars denote standard deviation. Tau factors were compared with an unpaired, two-sided student t-test. **(F)** Protein purity was assessed with a Coomassie stained SDS-PAGE. 5 μM of protein was loaded into each lane. **(G)** Western blot confirming protein expression of disease variants in transfected HEK293T lysates. 600 ng of HA tagged full-length ALX4 cDNA was transfected into a single well of a 6-well plate. β-actin was used as a loading control. HA (Roche: 3F10) staining is shown in red and β-actin (Licor: 926-42212) is shown in green.

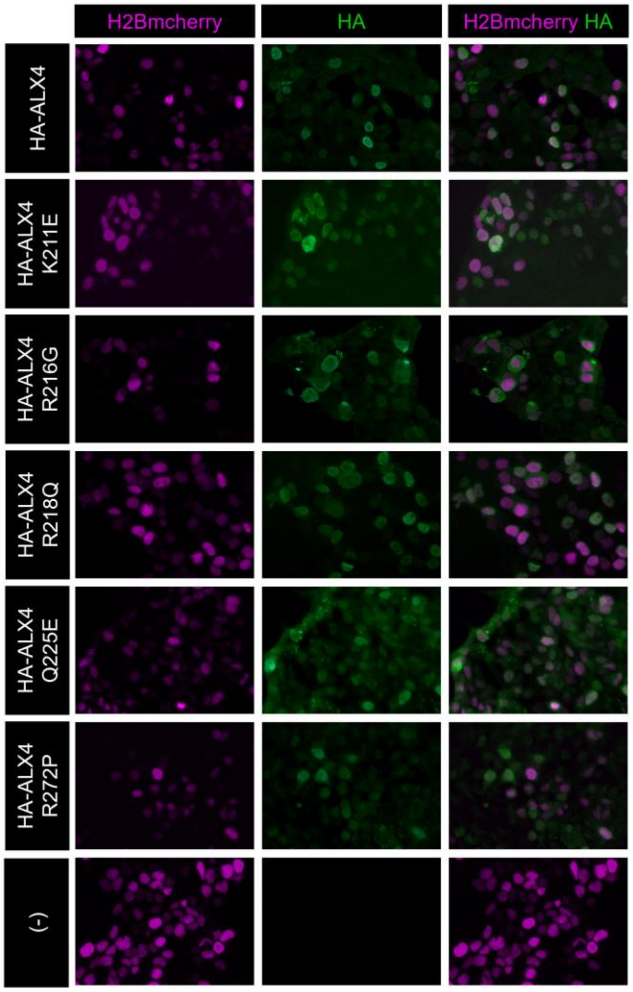


Supplementary Figure 9. Nuclear localization of ALX4 disease variants in transfected HEK293T cells. 40 ng of H2B-mcherry and 40 ng of corresponding HA tagged ALX4 cDNA in pcDNA3.1 were transfected into HEK293T cells. H2B-mcherry was used as a transfection control and its autofluorescence is shown in magenta. HA (Roche: 3F10) staining is shown in green. Note: brightness and contrast were individually adjusted to best match H2B and HA expression levels to determine localization. Expression levels cannot be gleaned from these images and were instead measured via western blot in Supplementary Figure 8G.

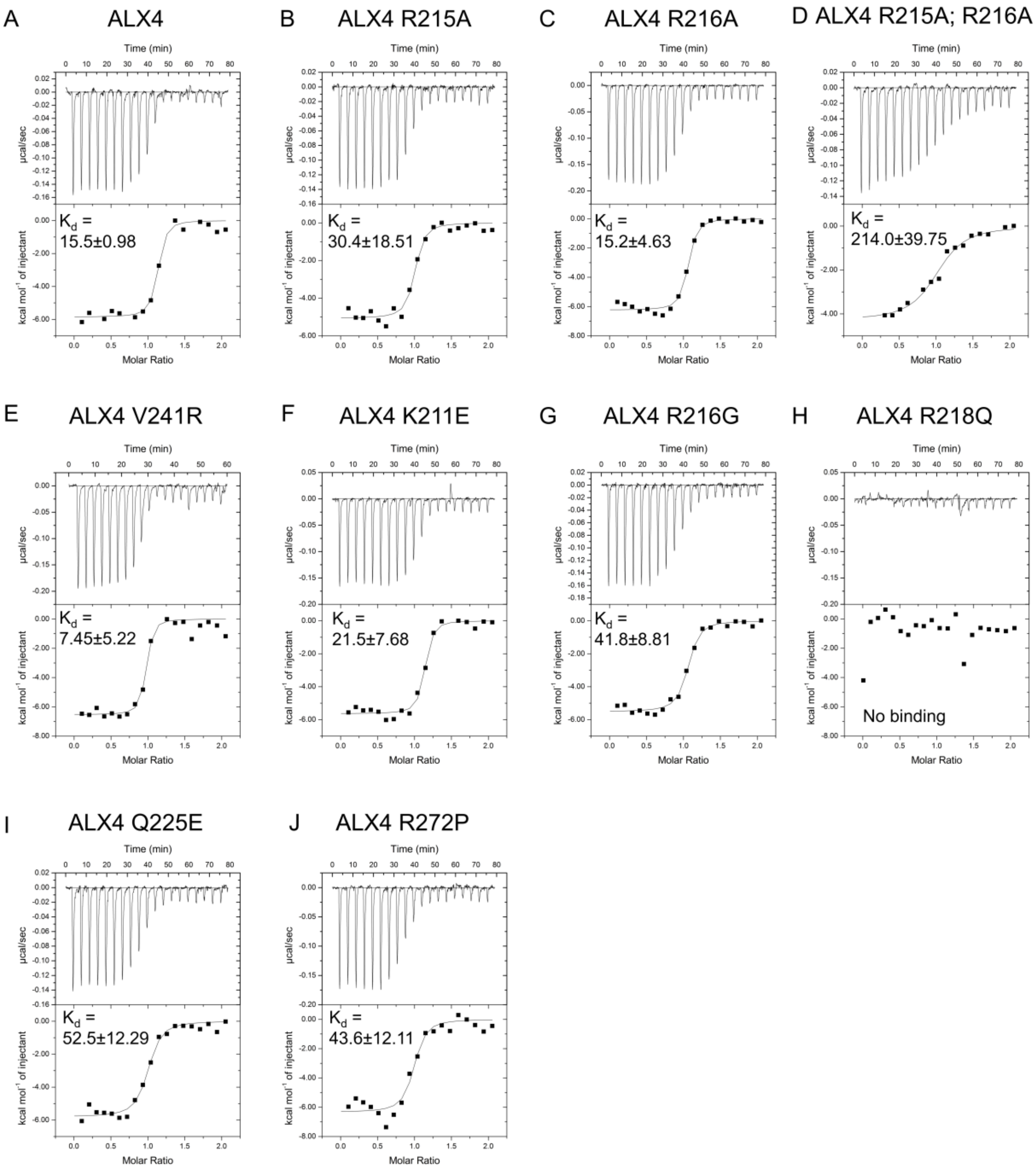


Supplementary Figure 10. Representative thermograms of heat signal for ALX4 structural and disease variants binding to the 14mer oligo duplex: CGCTAATTAGCTCG. The calculated dissociation constants ($K_d$) are printed on each plot in nM (Mean ± standard deviation).

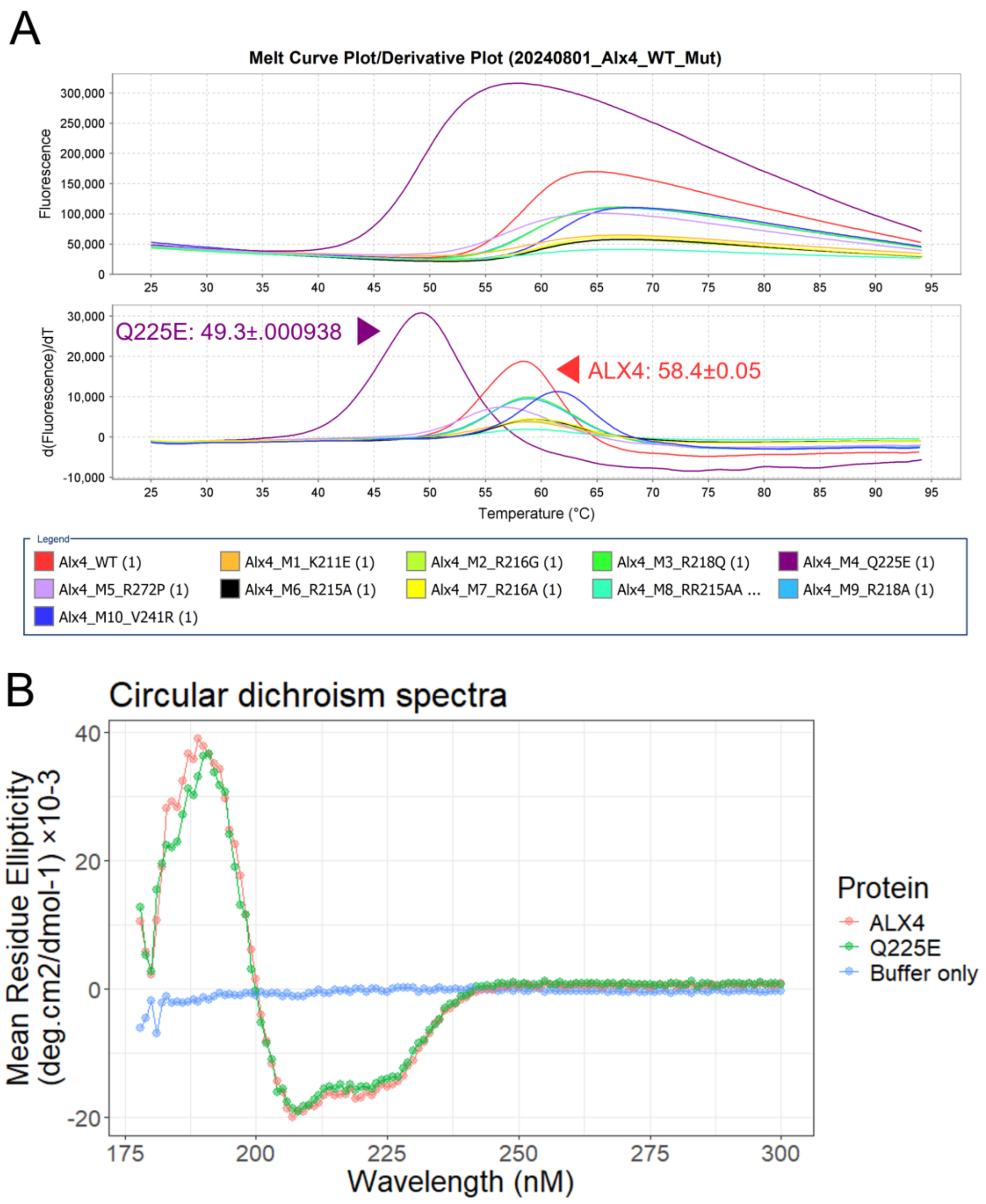


Supplementary Figure 11. Protein stability measurements of ALX4 structural mutations and ALX4 disease variants. **(A)** Melt curve plots from thermoshift testing of ALX4 structural and disease variants. Q225E is less stable than the wildtype ALX4 protein. **(B)** Circular dichroism spectra for ALX4 WT and ALX4 Q225E. There are no substantial differences between the secondary structure compositions of ALX4 WT and ALX4 Q225E.

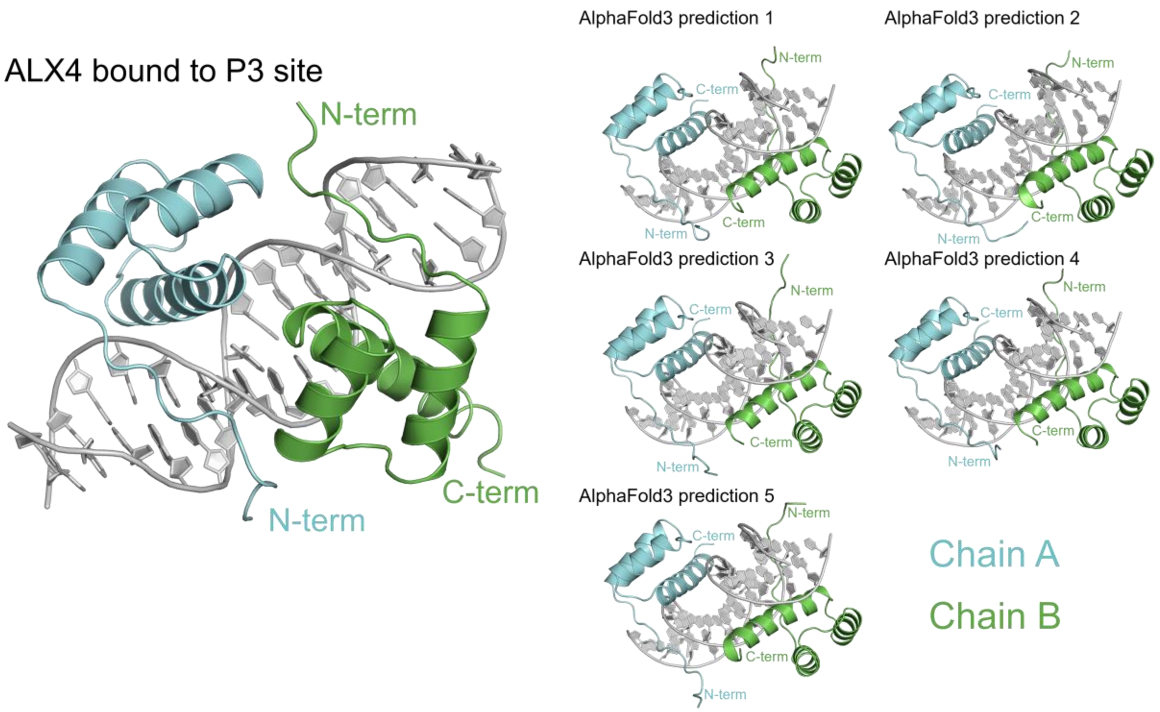


Supplementary Figure 12. AlphaFold3 fails to predict ALX4 dimer complex on the P3 site. ALX4 binds to the P3 site in a head-to-head orientation, however AlphaFold3 docks the two ALX4 proteins in a head-to-tail orientation for all predictions. AlphaFold3 failed to predict the three protein-protein interfaces as well as the key interacting residues.

**Supplementary information**

*Primers used for to generate cDNAs for protein purification*

> mAlx4 FWD primer

GGAgcggccgcaTCTGAACCCGTGGGCATGG

> mAlx4 REV primer

GGTctcgag**TTA**tttttcgaactgcgggtggctccaAATCTGGGCGTAGTTCTCAGC

> mAlx4 R215A FWD primer

taacaaaggcaagaaggcgcggaaccgaaccacc

> mAlx4 R215A REV primer

ggtggttcggttccgcgccttcttgcctttgtta

> mAlx4 R216A FWD primer

caaaggcaagaagcgggcgaaccgaaccaccttc

> mAlx4 R216A REV primer

gaaggtggttcggttcgcccgcttcttgcctttg

> mAlx4 R215A; R216A FWD primer

gagtaacaaaggcaagaaggcggcgaaccgaaccaccttcact

> mAlx4 R215A; R216A REV primer

agtgaaggtggttcggttcgccgccttcttgcctttgttactc

>mAlx4 V241R FWD primer

gctgctcccgtgcatacctatcagggtagtgtgtct

> mAlx4 V241R REV primer

agacacactaccctgataggtatgcacgggagcagc

> mAlx4 K211E FWD primer

gactcagagagcaacgagggcaagaagcggc

> mAlx4 K211E REV primer

gccgcttcttgccctcgttgctctctgagtc

> mAlx4 R216G FWD primer

ggcaagaagcgggggaaccggacca

> mAlx4 R216G REV primer

tggtccggttcccccgcttcttgcc

> mAlx4 R218Q FWD primer

aagcggcggaaccagaccaccttcacc

> mAlx4 R218Q REV primer

ggtgaaggtggtctggttccgccgctt

> mAlx4 Q225E FWD primer

ccttcaccagctacgagctggaggagctg

> mAlx4 Q225E REV primer

cagctcctccagctcgtagctggtgaagg

> mAlx4 R272P FWD primer

caagtggaggaagccggagcgttttgggc

> mAlx4 R272P REV primer

gcccaaaacgctccggcttcctccacttg

*EMSA probes*

> P3 site EMSA probe

TGTGTCTTAATTAGATTAACGCACTTGATAGTGCGGGCGTGGCT

> P4 site EMSA probe

TGTGTCTTAATTAG**G**ATTAACGCACTTGATAGTGCGGGCGTGGCT

> Fluorescently tagged probe

/5IRD700/AGCCACGCCCGCACTA

*Reporter sequences used in luciferase assays in pGL3 backbone*

5xUAS sequence ALX4 binding site Minimal promoter Restriction enzyme site

> 5xUAS-3xP3-minimal promoter

GGTACCTGAGCTCCTGCAGGTCGGAGTACTGTCCTCCGAGCGGAGTACTGTCCTCCGAGCGGAGTACTGTCCTCCGAGCGGAGTACTGTCCTCCGAGCGGAGTACTGTCCTCCGAGGGATCCGCTAGCCTCGAGTGTGTCTTAATTAGATTAACGCACTTGATGTGTCTTAATTAGATTAACGCACTTGATGTGTCTTAATTAGATTAACGCACTTGAAGATCTGGCCTCGGCGGCCAAGCTTAGACACTAGAGGGTATATAATGGAAGCTCGACTTCCAGCTTGGCAATCCGGTACTGTTGGTAAAGCCCCATGG

> 5xUAS-3xP4-minimal promoter

GGTACCTGAGCTCCTGCAGGTCGGAGTACTGTCCTCCGAGCGGAGTACTGTCCTCCGAGCGGAGTACTGTCCTCCGAGCGGAGTACTGTCCTCCGAGCGGAGTACTGTCCTCCGAGGGATCCGCTAGCCTCGAGTGTGTCTTAATTAG**G**ATTAACGCACTTGATGTGTCTTAATTAG**G**ATTAACGCACTTGATGTGTCTTAATTAG**G**ATTAACGCACTTGAAGATCTGGCCTCGGCGGCCAAGCTTAGACACTAGAGGGTATATAATGGAAGCTCGACTTCCAGCTTGGCAATCCGGTACTGTTGGTAAAGCCCCATGG

*Full length cDNAs of ALX4 variants*

HA tag Glycine linker Mutation

> HA-ALX4 in pcDNA3.1

ATGTACCCATACGATGTTCCAGATTACGCTGGCTCTGGAAATGCTGAGACTTGCGTCTCTTACTGCGAGTCGCCGGCCGCTGCCATGGACGCCTACTACAGCCCGGTGTCGCAGAGTCGGGAGGGCTCGTCGCCTTTTAGGGCATTTCCCGGAGGCGACAAGTTCGGCACAACTTTCCTGTCGGCCGCCGCCAAAGCACAGGGATTCGGGGACGCCAAGAGCCGGGCCCGTTACGGCGCTGGGCAGCAGGACCTGGCGACACCCCTGGAGAGTGGAGCTGGGGCGCGGGGCTCCTTTAACAAGTTCCAGCCCCAGCCGTCGACCCCGCAGCCCCAGCCGCCGCCGCAGCCGCAGCCGCAGCAGCAGCAGCCGCAGCCCCAGCCGCCCGCGCAACCGCATCTTTACTTGCAGCGAGGCGCCTGCAAGACGCCCCCGGACGGCAGCCTCAAACTCCAGGAAGGCAGCAGCGGCCACAGCGCGGCCTTGCAGGTTCCCTGCTACGCTAAAGAGAGCTCCCTGGGTGAGCCAGAGTTACCCCCTGACTCTGACACTGTGGGGATGGACAGCAGCTACCTGAGTGTCAAGGAGGCTGGGGTGAAGGGGCCCCAGGACCGGGCCAGCTCAGACCTCCCCAGCCCATTGGAGAAGGCCGACTCAGAGAGCAACAAGGGCAAGAAGCGGCGGAACCGGACCACCTTCACCAGCTACCAGCTGGAGGAGCTGGAGAAGGTCTTCCAGAAGACCCACTACCCAGACGTGTATGCGCGGGAACAGCTGGCCATGAGGACAGACCTCACTGAGGCCCGCGTGCAGGTCTGGTTCCAGAACCGAAGGGCCAAGTGGAGGAAGCGGGAGCGTTTTGGGCAGATGCAGCAGGTTCGAACCCACTTCTCCACTGCATATGAGCTGCCCCTCCTCACCCGAGCTGAGAACTACGCCCAGATTCAGAACCCGTCCTGGCTCGGCAACAACGGGGCTGCCTCACCAGTGCCAGCCTGCGTGGTCCCCTGCGACCCGGTGCCTGCCTGCATGTCCCCTCATGCCCACCCCCCTGGCTCTGGGGCCAGCAGCGTCACCGACTTCCTGAGTGTGTCTGGGGCTGGCAGTCACGTGGGCCAGACGCACATGGGCA

GCCTGTTTGGAGCAGCCAGCCTCAGCCCAGGCCTCAATGGCTACGAGCTCAACGGCGAGCCGGACCGCAAGACCTCGAGCATCGCGGCCCTCCGCATGAAGGCCAAGGAGCACAGTGCGGCCATTTCCTGGGCCACATGA

> HA-ALX4 V241R in pcDNA3.1

ATGTACCCATACGATGTTCCAGATTACGCTGGCTCTGGAAATGCTGAGACTTGCGTCTCTTACTGCGAGTCGCCGGCCGCTGCCATGGACGCCTACTACAGCCCGGTGTCGCAGAGTCGGGAGGGCTCGTCGCCTTTTAGGGCATTTCCCGGAGGCGACAAGTTCGGCACAACTTTCCTGTCGGCCGCCGCCAAAGCACAGGGATTCGGGGACGCCAAGAGCCGGGCCCGTTACGGCGCTGGGCAGCAGGACCTGGCGACACCCCTGGAGAGTGGAGCTGGGGCGCGGGGCTCCTTTAACAAGTTCCAGCCCCAGCCGTCGACCCCGCAGCCCCAGCCGCCGCCGCAGCCGCAGCCGCAGCAGCAGCAGCCGCAGCCCCAGCCGCCCGCGCAACCGCATCTTTACTTGCAGCGAGGCGCCTGCAAGACGCCCCCGGACGGCAGCCTCAAACTCCAGGAAGGCAGCAGCGGCCACAGCGCGGCCTTGCAGGTTCCCTGCTACGCTAAAGAGAGCTCCCTGGGTGAGCCAGAGTTACCCCCTGACTCTGACACTGTGGGGATGGACAGCAGCTACCTGAGTGTCAAGGAGGCTGGGGTGAAGGGGCCCCAGGACCGGGCCAGCTCAGACCTCCCCAGCCCATTGGAGAAGGCCGACTCAGAGAGCAACAAGGGCAAGAAGCGGCGGAACCGGACCACCTTCACCAGCTACCAGCTGGAGGAGCTGGAGAAGGTCTTCCAGAAGACCCACTACCCAGACAGGTATGCGCGGGAACAGCTGGCCATGAGGACAGACCTCACTGAGGCCCGCGTGCAGGTCTGGTTCCAGAACCGAAGGGCCAAGTGGAGGAAGCGGGAGCGTTTTGGGCAGATGCAGCAGGTTCGAACCCACTTCTCCACTGCATATGAGCTGCCCCTCCTCACCCGAGCTGAGAACTACGCCCAGATTCAGAACCCGTCCTGGCTCGGCAACAACGGGGCTGCCTCACCAGTGCCAGCCTGCGTGGTCCCCTGCGACCCGGTGCCTGCCTGCATGTCCCCTCATGCCCACCCCCCTGGCTCTGGGGCCAGCAGCGTCACCGACTTCCTGAGTGTGTCTGGGGCTGGCAGTCACGTGGGCCAGACGCACATGGGCAGCCTGTTTGGAGCAGCCAGCCTCAGCCCAGGCCTCAATGGCTACGAGCTCAACGGCGAGCCGGACCGCAAGACCTCGAGCATCGCGGCCCTCCGCATGAAGGCCAAGGAGCACAGTGCGGCCATTTCCTGGGCCACATGA

> HA-ALX4 K211E in pcDNA3.1

ATGTACCCATACGATGTTCCAGATTACGCTGGCTCTGGAAATGCTGAGACTTGCGTCTCTTACTGCGAGTCGCCGGCCGCTGCCATGGACGCCTACTACAGCCCGGTGTCGCAGAGTCGGGAGGGCTCGTCGCCTTTTAGGGCATTTCCCGGAG

GCGACAAGTTCGGCACAACTTTCCTGTCGGCCGCCGCCAAAGCACAGGGATTCGGGGACGCCAAGAGCCGGGCCCGTTACGGCGCTGGGCAGCAGGACCTGGCGACACCCCTGGAGAGTGGAGCTGGGGCGCGGGGCTCCTTTAACAAGTTCCAGCCCCAGCCGTCGACCCCGCAGCCCCAGCCGCCGCCGCAGCCGCAGCCGCAGCAGCAGCAGCCGCAGCCCCAGCCGCCCGCGCAACCGCATCTTTACTTGCAGCGAGGCGCCTGCAAGACGCCCCCGGACGGCAGCCTCAAACTCCAGGAAGGCAGCAGCGGCCACAGCGCGGCCTTGCAGGTTCCCTGCTACGCTAAAGAGAGCTCCCTGGGTGAGCCAGAGTTACCCCCTGACTCTGACACTGTGGGGATGGACAGCAGCTACCTGAGTGTCAAGGAGGCTGGGGTGAAGGGGCCCCAGGACCGGGCCAGCTCAGACCTCCCCAGCCCATTGGAGAAGGCCGACTCAGAGAGCAACGAGGGCAAGAAGCGGCGGAACCGGACCACCTTCACCAGCTACCAGCTGGAGGAGCTGGAGAAGGTCTTCCAGAAGACCCACTACCCAGACGTGTATGCGCGGGAACAGCTGGCCATGAGGACAGACCTCACTGAGGCCCGCGTGCAGGTCTGGTTCCAGAACCGAAGGGCCAAGTGGAGGAAGCGGGAGCGTTTTGGGCAGATGCAGCAGGTTCGAACCCACTTCTCCACTGCATATGAGCTGCCCCTCCTCACCCGAGCTGAGAACTACGCCCAGATTCAGAACCCGTCCTGGCTCGGCAACAACGGGGCTGCCTCACCAGTGCCAGCCTGCGTGGTCCCCTGCGACCCGGTGCCTGCCTGCATGTCCCCTCATGCCCACCCCCCTGGCTCTGGGGCCAGCAGCGTCACCGACTTCCTGAGTGTGTCTGGGGCTGGCAGTCACGTGGGCCAGACGCACATGGGCAGCCTGTTTGGAGCAGCCAGCCTCAGCCCAGGCCTCAATGGCTACGAGCTCAACGGCGAGCCGGACCGCAAGACCTCGAGCATCGCGGCCCTCCGCATGAAGGCCAAGGAGCACAGTGCGGCCATTTCCTGGGCCACATGA

> HA-ALX4 R216G in pcDNA3.1

ATGTACCCATACGATGTTCCAGATTACGCTGGCTCTGGAAATGCTGAGACTTGCGTCTCTTACTGCGAGTCGCCGGCCGCTGCCATGGACGCCTACTACAGCCCGGTGTCGCAGAGTCGGGAGGGCTCGTCGCCTTTTAGGGCATTTCCCGGAGGCGACAAGTTCGGCACAACTTTCCTGTCGGCCGCCGCCAAAGCACAGGGATTCGGGGACGCCAAGAGCCGGGCCCGTTACGGCGCTGGGCAGCAGGACCTGGCGACACCCCTGGAGAGTGGAGCTGGGGCGCGGGGCTCCTTTAACAAGTTCCAGCCCCAGCCGTCGACCCCGCAGCCCCAGCCGCCGCCGCAGCCGCAGCCGCAGCAGCAGCAGCCGCAGCCCCAGCCGCCCGCGCAACCGCATCTTTACTTGCAGCGAGGCGCCTGCAAGACGCCCCCGGACGGCAGCCTCAAACTCCAGGAAGGCAGCAGCGGCCACAGCGCGGCCTTGCAGGTTCCCTGCTACGCTAAAGAGAGCTCCCTGGGTGAGCCAGAGTTACCCCCTGACTCTGACACTGTGGGGATGGACAGCAGCTACCTGAGTGTCAAGGAGGCTGGGGTGAAGGGGC

CCCAGGACCGGGCCAGCTCAGACCTCCCCAGCCCATTGGAGAAGGCCGACTCAGAGAGCAACAAGGGCAAGAAGCGGGGGAACCGGACCACCTTCACCAGCTACCAGCTGGAGGAGCTGGAGAAGGTCTTCCAGAAGACCCACTACCCAGACGTGTATGCGCGGGAACAGCTGGCCATGAGGACAGACCTCACTGAGGCCCGCGTGCAGGTCTGGTTCCAGAACCGAAGGGCCAAGTGGAGGAAGCGGGAGCGTTTTGGGCAGATGCAGCAGGTTCGAACCCACTTCTCCACTGCATATGAGCTGCCCCTCCTCACCCGAGCTGAGAACTACGCCCAGATTCAGAACCCGTCCTGGCTCGGCAACAACGGGGCTGCCTCACCAGTGCCAGCCTGCGTGGTCCCCTGCGACCCGGTGCCTGCCTGCATGTCCCCTCATGCCCACCCCCCTGGCTCTGGGGCCAGCAGCGTCACCGACTTCCTGAGTGTGTCTGGGGCTGGCAGTCACGTGGGCCAGACGCACATGGGCAGCCTGTTTGGAGCAGCCAGCCTCAGCCCAGGCCTCAATGGCTACGAGCTCAACGGCGAGCCGGACCGCAAGACCTCGAGCATCGCGGCCCTCCGCATGAAGGCCAAGGAGCACAGTGCGGCCATTTCCTGGGCCACATGA

> HA-ALX4 R218Q in pcDNA3.1

ATGTACCCATACGATGTTCCAGATTACGCTGGCTCTGGAAATGCTGAGACTTGCGTCTCTTACTGCGAGTCGCCGGCCGCTGCCATGGACGCCTACTACAGCCCGGTGTCGCAGAGTCGGGAGGGCTCGTCGCCTTTTAGGGCATTTCCCGGAGGCGACAAGTTCGGCACAACTTTCCTGTCGGCCGCCGCCAAAGCACAGGGATTCGGGGACGCCAAGAGCCGGGCCCGTTACGGCGCTGGGCAGCAGGACCTGGCGACACCCCTGGAGAGTGGAGCTGGGGCGCGGGGCTCCTTTAACAAGTTCCAGCCCCAGCCGTCGACCCCGCAGCCCCAGCCGCCGCCGCAGCCGCAGCCGCAGCAGCAGCAGCCGCAGCCCCAGCCGCCCGCGCAACCGCATCTTTACTTGCAGCGAGGCGCCTGCAAGACGCCCCCGGACGGCAGCCTCAAACTCCAGGAAGGCAGCAGCGGCCACAGCGCGGCCTTGCAGGTTCCCTGCTACGCTAAAGAGAGCTCCCTGGGTGAGCCAGAGTTACCCCCTGACTCTGACACTGTGGGGATGGACAGCAGCTACCTGAGTGTCAAGGAGGCTGGGGTGAAGGGGCCCCAGGACCGGGCCAGCTCAGACCTCCCCAGCCCATTGGAGAAGGCCGACTCAGAGAGCAACAAGGGCAAGAAGCGGCGGAACCAGACCACCTTCACCAGCTACCAGCTGGAGGAGCTGGAGAAGGTCTTCCAGAAGACCCACTACCCAGACGTGTATGCGCGGGAACAGCTGGCCATGAGGACAGACCTCACTGAGGCCCGCGTGCAGGTCTGGTTCCAGAACCGAAGGGCCAAGTGGAGGAAGCGGGAGCGTTTTGGGCAGATGCAGCAGGTTCGAACCCACTTCTCCACTGCATATGAGCTGCCCCTCCTCACCCGAGCTGAGAACTACGCCCAGATTCAGAACCCGTCCTGGCTCGGCAACAACGGGGCTGCCTCACCAGTGCCAGCCTGCGTGGTCCCCTGCGACCCGGTGCCTGCCTGCATGTCCCCTCATGCCCACCCCCCTGGCT

CTGGGGCCAGCAGCGTCACCGACTTCCTGAGTGTGTCTGGGGCTGGCAGTCACGTGGGCCAGACGCACATGGGCAGCCTGTTTGGAGCAGCCAGCCTCAGCCCAGGCCTCAATGGCTACGAGCTCAACGGCGAGCCGGACCGCAAGACCTCGAGCATCGCGGCCCTCCGCATGAAGGCCAAGGAGCACAGTGCGGCCATTTCCTGGGCCACATGA

> HA-ALX4 Q225E in pcDNA3.1

ATGTACCCATACGATGTTCCAGATTACGCTGGCTCTGGAAATGCTGAGACTTGCGTCTCTTACTGCGAGTCGCCGGCCGCTGCCATGGACGCCTACTACAGCCCGGTGTCGCAGAGTCGGGAGGGCTCGTCGCCTTTTAGGGCATTTCCCGGAGGCGACAAGTTCGGCACAACTTTCCTGTCGGCCGCCGCCAAAGCACAGGGATTCGGGGACGCCAAGAGCCGGGCCCGTTACGGCGCTGGGCAGCAGGACCTGGCGACACCCCTGGAGAGTGGAGCTGGGGCGCGGGGCTCCTTTAACAAGTTCCAGCCCCAGCCGTCGACCCCGCAGCCCCAGCCGCCGCCGCAGCCGCAGCCGCAGCAGCAGCAGCCGCAGCCCCAGCCGCCCGCGCAACCGCATCTTTACTTGCAGCGAGGCGCCTGCAAGACGCCCCCGGACGGCAGCCTCAAACTCCAGGAAGGCAGCAGCGGCCACAGCGCGGCCTTGCAGGTTCCCTGCTACGCTAAAGAGAGCTCCCTGGGTGAGCCAGAGTTACCCCCTGACTCTGACACTGTGGGGATGGACAGCAGCTACCTGAGTGTCAAGGAGGCTGGGGTGAAGGGGCCCCAGGACCGGGCCAGCTCAGACCTCCCCAGCCCATTGGAGAAGGCCGACTCAGAGAGCAACAAGGGCAAGAAGCGGCGGAACCGGACCACCTTCACCAGCTACGAGCTGGAGGAGCTGGAGAAGGTCTTCCAGAAGACCCACTACCCAGACGTGTATGCGCGGGAACAGCTGGCCATGAGGACAGACCTCACTGAGGCCCGCGTGCAGGTCTGGTTCCAGAACCGAAGGGCCAAGTGGAGGAAGCGGGAGCGTTTTGGGCAGATGCAGCAGGTTCGAACCCACTTCTCCACTGCATATGAGCTGCCCCTCCTCACCCGAGCTGAGAACTACGCCCAGATTCAGAACCCGTCCTGGCTCGGCAACAACGGGGCTGCCTCACCAGTGCCAGCCTGCGTGGTCCCCTGCGACCCGGTGCCTGCCTGCATGTCCCCTCATGCCCACCCCCCTGGCTCTGGGGCCAGCAGCGTCACCGACTTCCTGAGTGTGTCTGGGGCTGGCAGTCACGTGGGCCAGACGCACATGGGCAGCCTGTTTGGAGCAGCCAGCCTCAGCCCAGGCCTCAATGGCTACGAGCTCAACGGCGAGCCGGACCGCAAGACCTCGAGCATCGCGGCCCTCCGCATGAAGGCCAAGGAGCACAGTGCGGCCATTTCCTGGGCCACATGA

> HA-ALX4 R272P in pcDNA3.1

ATGTACCCATACGATGTTCCAGATTACGCTGGCTCTGGAAATGCTGAGACTTGCGTCTCTTACTGCGAGTCGCCGGCCGCTGCCATGGACGCCTACTACAGCCCGGTGTCGCAGAGTCGGGAGGGCTCGTCGCCTTTTAGGGCATTTCCCGGAGGCGACAAGTTCGGCACAACTTTCCTGTCGGCCGCCGCCAAAGCACAGGGATTCGGGGACGCCAAGAGCCGGGCCCGTTACGGCGCTGGGCAGCAGGACCTGGCGACACCCCTGGAGAGTGGAGCTGGGGCGCGGGGCTCCTTTAACAAGTTCCAGCCCCAGCCGTCGACCCCGCAGCCCCAGCCGCCGCCGCAGCCGCAGCCGCAGCAGCAGCAGCCGCAGCCCCAGCCGCCCGCGCAACCGCATCTTTACTTGCAGCGAGGCGCCTGCAAGACGCCCCCGGACGGCAGCCTCAAACTCCAGGAAGGCAGCAGCGGCCACAGCGCGGCCTTGCAGGTTCCCTGCTACGCTAAAGAGAGCTCCCTGGGTGAGCCAGAGTTACCCCCTGACTCTGACACTGTGGGGATGGACAGCAGCTACCTGAGTGTCAAGGAGGCTGGGGTGAAGGGGCCCCAGGACCGGGCCAGCTCAGACCTCCCCAGCCCATTGGAGAAGGCCGACTCAGAGAGCAACAAGGGCAAGAAGCGGCGGAACCGGACCACCTTCACCAGCTACCAGCTGGAGGAGCTGGAGAAGGTCTTCCAGAAGACCCACTACCCAGACGTGTATGCGCGGGAACAGCTGGCCATGAGGACAGACCTCACTGAGGCCCGCGTGCAGGTCTGGTTCCAGAACCGAAGGGCCAAGTGGAGGAAGCCGGAGCGTTTTGGGCAGATGCAGCAGGTTCGAACCCACTTCTCCACTGCATATGAGCTGCCCCTCCTCACCCGAGCTGAGAACTACGCCCAGATTCAGAACCCGTCCTGGCTCGGCAACAACGGGGCTGCCTCACCAGTGCCAGCCTGCGTGGTCCCCTGCGACCCGGTGCCTGCCTGCATGTCCCCTCATGCCCACCCCCCTGGCTCTGGGGCCAGCAGCGTCACCGACTTCCTGAGTGTGTCTGGGGCTGGCAGTCACGTGGGCCAGACGCACATGGGCAGCCTGTTTGGAGCAGCCAGCCTCAGCCCAGGCCTCAATGGCTACGAGCTCAACGGCGAGCCGGACCGCAAGACCTCGAGCATCGCGGCCCTCCGCATGAAGGCCAAGGAGCACAGTGCGGCCATTTCCTGGGCCACATGA

Supplementary table 1. X-ray Data collection and refinement statistics.

**Data Collection Statistics**

| Complex | Alx4.HD-DNA | Alx4.HD |
|---|---|---|
| Resolution (Å) | 79.57 – 2.39 (2.48 – 2.39) | 67.73-2.39 (2.48 – 2.39) |
| Space Group | $P3_221$ | I23 |
| Wavelength (Å) | 0.97872 | 0.97872 |
| Unit Cell a, b, c (Å) | 70.45, 70.45, 159.14 | 95.78, 95.78, 95.78 |
| Unit Cell $\alpha$, $\beta$, $\gamma$ (°) | 90.00, 90.00, 120.00 | 90.00, 90.00, 90.00 |
| $R_{merge}$ | 0.146 (3.245) | 0.118 (2.752) |
| $I/\sigma I$ | 4.7 (0.6) | 14.1 (1.3) |
| $CC_{1/2}$ | 0.987 (0.638) | 0.999 (0.495) |
| Completeness (%) | 100.0 (100.0) | 100.0 (100.0) |
| Redundancy | 10.7 (9.7) | 21.9 (19.8) |

**Refinement Statistics**

| | | |
|---|---|---|
| $R_{work}/R_{free}$ (%) | 21.36 / 23.88 (37.74 / 49.73) | 21.41 / 24.86 (34.60 / 37.84) |
| Number of reflections | 18,810 | 5,952 |
| Number of atoms | 1,881 | 578 |
| Complexes/asymmetric unit | 1 | 1 |
| Wilson B/Mean B value (Å$^2$) | 95.71 / 122.87 | 68.01 |
| RMSD Bond Lengths (Å) | 0.008 | 0.008 |
| RMSD Bond Angles (°) | 0.82 | 0.992 |
| Ramachandran (favored/outliers) | 95.38% / 0.00% | 100.0% / 0.00% |

Highest resolution shell shown in parentheses.

Supplementary table 2. Gene expression omnibus ID for analyzed datasets

| Assay type | Pulldown | TWIST1 depleted? | Replicate | GSM ID | SRR ID | Cell line |
|---|---|---|---|---|---|---|
| CUT&RUN | ALX4 | | 1 | GSM7213748 | SRR24258480 | TWIST1FV |
| CUT&RUN | TWIST1 | | 1 | GSM7213747 | SRR24258481 | TWIST1FV |
| CUT&RUN | IgG | | 1 | GSM7213753 | SRR24258475 | TWIST1FV |
| CUT&RUN | ALX4 | YES | 1 | GSM7213755 | SRR24258473 | TWIST1FV |
| CUT&RUN | IgG | YES | 1 | GSM7213760 | SRR24258468 | TWIST1FV |
| ChIP | H3K27ac | | 1 | GSM7213607 | SRR24257082 | TWIST1FV |
| ChIP | H3K27ac | | 2 | GSM7213610 | SRR24257081 | TWIST1FV |
| ChIP | IgG | | 1 | GSM7213617 | SRR24257051 | TWIST1FV |
| ChIP | IgG | | 2 | GSM7213620 | SRR24257016 | TWIST1FV |
| ATAC | | | 1 | GSM7213525 | SRR24257841 | TWIST1FV |
| ATAC | | | 2 | GSM7213528 | SRR24257838 | TWIST1FV |
| ATAC | | YES | 1 | GSM7213526 | SRR24257840 | TWIST1FV |
| ATAC | | YES | 2 | GSM7213529 | SRR24257837 | TWIST1FV |
